# Cell polarity proteins promote macropinocytosis in response to metabolic stress

**DOI:** 10.1101/2024.01.16.575943

**Authors:** Guillem Lambies, Szu-Wei Lee, Karen Duong-Polk, Pedro Aza-Blanc, Swetha Maganti, David W. Dawson, Cosimo Commisso

## Abstract

Macropinocytosis has emerged as a nutrient-scavenging pathway that cancer cells exploit to survive the nutrient-deprived conditions of the tumor microenvironment. Cancer cells are especially reliant on glutamine for their survival, and in pancreatic ductal adenocarcinoma (PDAC) cells, glutamine deficiency can enhance the stimulation of macropinocytosis, allowing the cells to escape metabolic stress through the production of extracellular-protein-derived amino acids. Here, we identify the atypical protein kinase C (aPKC) enzymes, PKCζ and PKCι, as novel regulators of macropinocytosis. In normal epithelial cells, aPKCs are known to regulate cell polarity in association with the scaffold proteins Par3 and Par6, controlling the function of several targets, including the Par1 kinases. In PDAC cells, we identify that each of these cell polarity proteins are required for glutamine stress-induced macropinocytosis. Mechanistically, we find that the aPKCs are regulated by EGFR signaling or by the transcription factor CREM to promote the relocation of Par3 to microtubules, facilitating macropinocytosis in a dynein-dependent manner. Importantly, we determine that cell fitness impairment caused by aPKC depletion is rescued by the restoration of macropinocytosis and that aPKCs support PDAC growth *in vivo*. These results identify a previously unappreciated role for cell polarity proteins in the regulation of macropinocytosis and provide a better understanding of the mechanistic underpinnings that control macropinocytic uptake in the context of metabolic stress.

## Introduction

Macropinocytosis, also known as “cellular drinking”, is a clathrin-independent endocytic pathway that non-selectively internalizes extracellular cargo into large vesicles known as macropinosomes^1–3^. In Ras-mutant cells, this uptake pathway is an amino acid supply route that allows the cancer cells to survive and proliferate despite the nutrient scarcity present in the tumor microenviroment^4–6^. By internalizing extracellular proteins, such as serum albumin, and targeting them for lysosome-dependent degradation, macropinocytosis provides a source of protein-derived amino acids. Under physiological conditions, these protein-derived amino acids can account for almost half of the intracellular and extracellular amino acid pools^7^.Importantly, it has been well established in multiple mouse models of cancer that macropinocytosis inhibition *in vivo* significantly suppresses tumor growth.

Macropinocytosis in Ras-mutant cells is elevated as a response to amino acid deprivation^3, 5, 6, 8^. The vast majority of pancreatic ductal adenocarcinoma (PDAC) tumors harbor a mutation in *KRAS,* and we previously demonstrated that macropinocytosis in PDAC cells is considerably enhanced when glutamine is limiting^5^. This is physiologically important since in human PDAC tumors glutamine is the most depleted amino acid relative to adjacent non-neoplastic tissue^7^. Glutamine is a critical nutrient in tumors as it supports various aspects of cancer metabolism including lipid and protein biosynthesis, the *de novo* synthesis of nucleotides, the urea cycle, glutathione production and the biosynthesis of other amino acids^9^. Downstream aspects of the macropinocytosis pathway are also impacted by amino acid deprivation. For example, in *KRAS*-mutant mouse embryonic fibroblasts (MEFs), the lysosome-dependent degradation of macropinocytosed protein is enhanced by amino acid starvation ^8^. This is consistent with the observation that in PDAC xenograft tumors, the highest macropinocytic rates are detected in the tumor core regions, which are depleted of glutamine and other amino acids relative to the tumor periphery^5^. While the functional importance of macropinocytosis in PDAC tumors is highly appreciated, the specific regulators that control macropinocytosis in the context of nutrient stress remain poorly understood.

Here, we identify PKCζ and PKCι, which constitute the family of atypical Protein Kinase C (aPKC) enzymes, as novel regulators of macropinocytosis. In normal epithelial cells, aPKCs function in maintaining cell polarity, a cellular process that is lost in cancer cells. However, we find that with glutamine stress, aPKCs are repurposed in PDAC cells, along with other cell polarity proteins such as Par-1a, Par3, and Par6, to drive macropinocytosis. In cell polarity, aPKCs control microtubule dynamics^10, 11^, which are essential to maintain epithelial cells in a polarized state^11, 12^. Interestingly, in PDAC cells, we find that the microtubule cytoskeleton and dynein-dependent microtubule transport are required for macropinocytosis, and that this is associated with aPKC-dependent redistribution of Par3 to microtubules. Mechanistically, we find that glutamine deficiency primes the aPKCs for activation by EGFR. Glutamine stress can also be triggered by specifically targeting glutamine-utilizing enzymes, as is the case with 6-diazo-5-oxo-L-norleucine (DON), a glutamine antagonist that enhances macropinocytosis in PDAC cells^5^. Interestingly, we find that pharmacological inhibition of glutamine metabolism with DON increases expression of the aPKCs through the transcription factor CREM. Functionally, we demonstrate that in low glutamine conditions, aPKC depletion leads to reduced proliferative capacity, a phenotype that is rescued when macropinocytosis is restored. Moreover, we elucidate that aPKCs regulate macropinocytosis *in vivo* and that they are essential to maintain PDAC tumor growth. Our interrogations of human PDAC clinical samples and datasets reveal that aPKCs are upregulated in tumors, contribute to patient survival, and, in a subset of tumors, are primed for activation. Altogether, our work sheds light on the mechanistic underpinnings that drive macropinocytosis in response to metabolic stress and point to the possibility of targeting cell polarity proteins in cancer.

## Results

### PKCζ and PKCι regulate macropinocytosis in PDAC cells

Metabolic stress caused by glutamine deprivation stimulates macropinocytosis; however, a comprehensive understanding of modulators that can control uptake in the context of nutrient stress is lacking^5^. In order to gain insight into the pathways that might broadly function in macropinocytosis in the context of glutamine stress, we performed a high-throughput (HT) siRNA screen using a HT-compatible macropinocytosis assay that we previously developed^13, 14^. For the screen, we used AsPC-1 cells since these cells display robust stimulation of macropinocytosis upon glutamine starvation (Extended Data Fig. 1a,b). Macropinocytosis was detected through the uptake of FITC-dextran, an established marker of macropinosomes and macropinosomes were quantified by image-based high-content analysis. Using our HT assay, we performed a kinome-wide siRNA screen (siOTP; Dharmacon) to identify kinases involved in controlling macropinocytosis in cells that were exposed to glutamine stress. With this approach, we identified the atypical Protein Kinase C (aPKC) isoforms PKCζ and PKCι as novel modulators of glutamine-stress-induced macropinocytosis (Fig. 1a). We next validated the role of the aPKCs in macropinocytosis by performing knockdown experiments using independent siRNAs from a different library, which allowed us to mitigate potential off-target effects. For validation, in addition to AsPC-1 cells, we also used HPAF-II cells since these cells display robust activation of macropinocytosis by glutamine stress (Extended Data Fig. 1c,d). Specific knockdown of the individual isoforms in AsPC-1 and HPAF-II cells was confirmed by western blot (Fig.1b,c). Validating a role for each aPKC in macropinocytosis, we found that knockdown of either PKCζ or PKCι suppressed macropinocytosis in glutamine-deprived conditions (Fig. 1d-g). In addition, we also observed a significant diminishment of macropinocytosis in the same conditions when cells were treated with ACPD, a specific pharmacological inhibitor of the aPKCs^15^(Fig. 1h-k). Altogether, our data indicate that the aPKCs are required to control macropinocytosis in the context of glutamine stress.

**Fig. 1.**
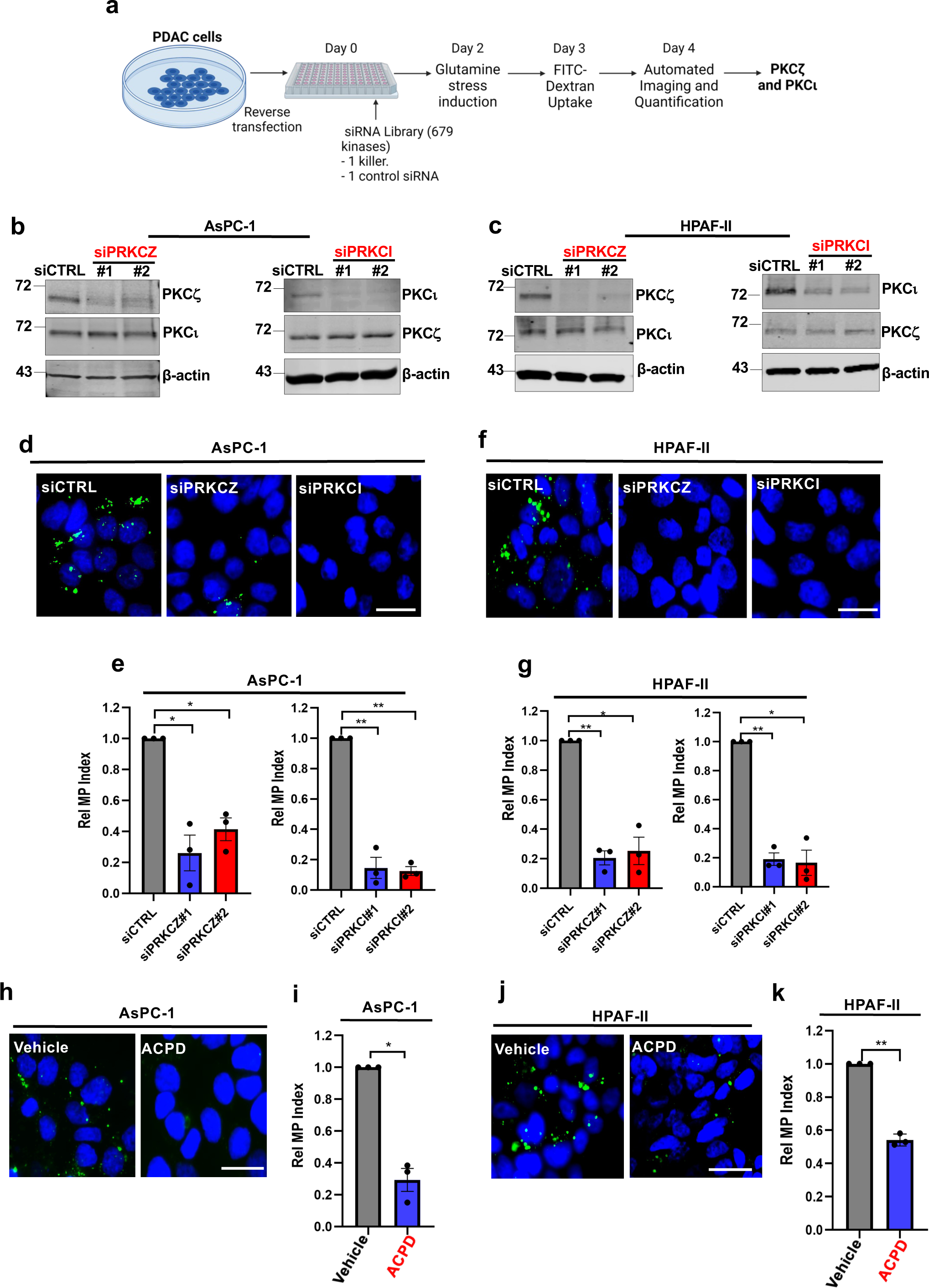
PKCζ and PKCι are required for glutamine stress-induced macropinocytosis in PDAC cells. **a)** Schematic depicting the steps employed for the siRNA kinome-wide screen in glutamine-stressed AsPC-1 cells. **b, c**) Immunoblots assessing PKCζ and PKCι levels in AsPC-1 (b) and HPAF-II (c) cells upon transfection with non-targeting control siRNA (siCTRL) or siRNAs targeting *PRKCZ* (siPRKCZ#1 and siPRKCZ#2) or *PRKCI* (siPRKCI#1 and siPRKCI#2). β-actin was used as a loading control. **d)** Representative fluorescent microscopy images from a FITC-dextran macropinocytosis assay (green) in glutamine-starved AsPC-1 cells transfected with the indicated siRNAs. Nuclei are stained with DAPI (blue). Scale bar, 20 μm. **e)** Quantification of macropinocytosis in glutamine-starved AsPC-1 cells transfected with the indicated siRNAs. Data are shown relative to siCTRL. Data are presented as mean ± SEM from three independent experiments. **f)** Representative images of macropinocytosis in glutamine-starved HPAF-II cells. Scale bar, 20 μm. **g)** Quantification of macropinocytosis in HPAF-II cells under the conditions described in f. Data are shown relative to siCTRL. Data are presented as mean ± SEM from three independent experiments. **h)** Representative images of macropinocytosis in glutamine-starved AsPC-1 cells treated with vehicle (water) or ACPD at 4μM for 72 hrs. Scale bar, 20 μm. **i)** Quantification of macropinocytosis for the conditions described in h. Data are shown relative to vehicle control. Data are presented as mean ± SEM from three independent experiments. **j)** Representative images of macropinocytosis in glutamine-starved HPAF-II cells treated with vehicle (water) or ACPD at 25 μM for 72 hrs. Scale bar, 20 μm. **k)** Quantification of macropinocytosis for the conditions described in j. Data are shown relative to vehicle control. Data are presented as mean ± SEM from three independent experiments. Statistical significance was calculated using unpaired two-tailed Student’s t test with Welch’s correction. \**P*<0.05, \*\**P*<0.01.

### PAR proteins and microtubule transport are required for glutamine stress-induced macropinocytosis

aPKCs are known to regulate apical-basal polarity in epithelial cells, which is essential for maintaining epithelial structure and integrity^16, 17^. In cancer, the prevailing notion is that cell polarity is lost; however, different members of this protein network are known to be involved in several tumorigenic processes^17–19^. In cell polarity, aPKCs associate with the scaffold proteins Par3 and Par6 constituting the Par or cell polarity complex that controls the function and subcellular localization of multiple cell polarity related substrates, such as Par1 kinases^10, 11, 17, 20–22^. In our screen, in addition to the aPKCs, we also identified the microtubule-associated Par-1a kinase as a ‘hit’. Considering the role of aPKCs and Par1 kinases in the modulation of epithelial cell polarity, we hypothesized that cell polarity proteins might be broadly required for regulating glutamine stress-driven macropinocytosis. To test this idea, we investigated the Par-1a kinase and the scaffold proteins Par3 and Par6 since these proteins form a complex with aPKC that regulates cell polarity. We found that knockdown of Par-1a, Par3, or Par6 suppressed macropinocytosis to a similar extent as aPKC depletion in glutamine-starved conditions (Fig. 2a,b; Extended Data Fig. 1e-h). These data highlight a novel and previously unrecognized role for cell polarity proteins in the regulation of macropinocytosis.

**Fig. 2.**
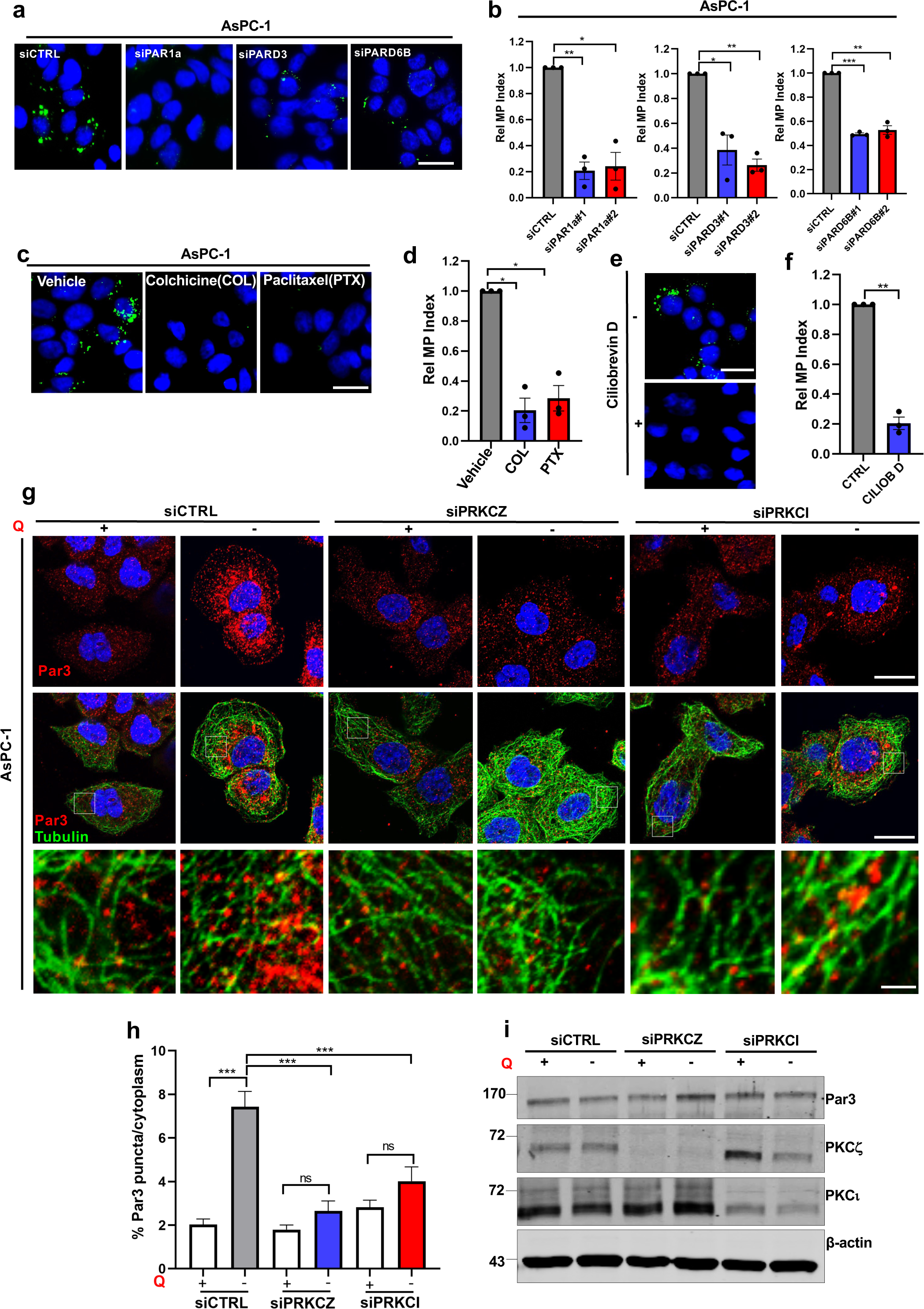
Par polarity proteins and the microtubule network are required for macropinocytosis. **a)** Representative images of macropinocytosis in glutamine-starved AsPC-1 cells transfected with siCTRL or siRNAs targeting *PAR1A* (siPAR1a), *PARD3* (siPARD3) and *PARD6B* (siPARD6B). Scale bar, 20 μm. **b)** Quantification of macropinocytosis in glutamine-starved AsPC-1 cells transfected with the indicated siRNAs. Data are shown relative to siCTRL. Data are presented as mean ± SEM from three independent experiments. **c)** Representative images of macropinocytosis in glutamine-starved AsPC-1 cells treated for 18 hrs with vehicle (water) or the microtubule inhibitors colchicine (COL, 100 ng/mL) and paclitaxel (PTX, 200 nM). Scale bar, 20μm. **d)** Quantification of macropinocytosis for the conditions described in c. Data are shown relative to vehicle control. Data are presented as mean ± SEM from three independent experiments. **e)** Representative images of macropinocytosis in glutamine-starved AsPC-1 cells treated with vehicle (water) or the dynein inhibitor ciliobrevin D (CILIOB D) for 30 minutes at 50 μM. Scale bar, 20μm. **f)** Quantification of macropinocytosis for the conditions described in e. Data are shown relative to vehicle control. Data are presented as mean ± SEM from three independent experiments. **g)** Representative images of Par3 protein (red) immunostaining in AsPC-1 cells transfected with the indicated siRNAs in glutamine-replete (+) or glutamine-free (−) media. Anti-Tubulin staining is shown in green. Nuclei are stained with DAPI (blue). Scale 20 μm. Bottom row are higher magnification images of the boxed areas. Scale bar, 5 μm. **h)** Quantification of the percent of Par3 protein found in subcellular puncta versus the cytoplasm. At least n=40 cells per condition were analyzed. Data are presented as mean ± SEM from three independent experiments. **i)** Immunoblot of Par3, PKCζ and PKCι proteins for the conditions described in (g). β-actin was used as a loading control. Data are representative of three independent experiments. Statistical significance was calculated using unpaired two-tailed Student’s t test with Welch’s correction. Ns, non-significant, \**P*<0.05, \*\**P*<0.01, \*\*\**P*<0.001.

In the establishment of cell polarity, there is specific spatiotemporal regulation of both actin and microtubule dynamics by the Par complex^12, 21^. While the role of actin in macropinocytosis has been extensively studied, it is not clear the extent to which microtubules contribute to macropinocytosis. There is some evidence suggesting that microtubules might be involved in the post-internalization trafficking events of mature macropinosomes^23^, but whether microtubules function in nascent macropinosome formation has not been examined. Since aPKCs are known to modulate microtubule dynamics^12^, and we detected the microtubule-associated Par-1a kinase as a macropinocytic modulator, we further examined whether microtubules could be involved in regulating macropinocytic uptake. We found that disruption of the microtubule network with either colchicine or paclitaxel, which would destabilize or stabilize microtubules, respectively, impaired glutamine stress-driven macropinocytosis (Fig. 2c,d). We next evaluated whether glutamine stress had the capacity to alter the morphology of the microtubule network. In control cells, glutamine deprivation caused the reorganization of the microtubules such that tubulin was more greatly associated with plasma membrane ruffles; however, this was not dependent on aPKCs (Extended Data Fig. 2a). These data suggested that aPKCs might be regulating macropinocytosis independent of plasma membrane ruffling. To form a nascent macropinosome, it is thought that cytoskeletal contractile forces are required to propel the macropinosome inwards, separating the macropinosome from the plasma membrane in a scission or fission process. To test this idea, we examined the function of the microtubule-associated motor protein dynein, which mediates the retrograde-directed transport of different cargoes along the microtubule network, including endocytic vesicles such as endosomes^24^. Dynein inhibition with ciliobrevin D nearly completely blocked macropinocytosis (Fig. 2e,f), suggesting that aPKCs could be mediating macropinosome formation through mechanisms involving the regulation of microtubule motor functions.

The cell polarity protein that is known to associate with dynein in the regulation of microtubule motor properties is the scaffold protein Par3^25, 26^. In the establishment and maintenance of cell polarity, Par3 subcellular localization is in part controlled by aPKCs that phosphorylate Par3 to promote its dissociation from the Par complex and its relocation to adherens junctions^27–29^. Considering that Par3 loss efficiently impaired macropinocytosis, we next examined whether glutamine stress was altering Par3 subcellular localization, and whether any observed changes in Par3 cellular distribution were dependent on aPKC. Remarkably, we found that glutamine deprivation promoted the localization of Par3 into microtubule-associated puncta, which was dependent on aPKCs (Fig. 2g,h; Extended Data Fig. 2b,c). We found that the enhancement of Par3 protein on the microtubules was due to a redistribution of cytosolic Par3 protein to the microtubules and not due to an increase in overall Par3 protein levels (Fig. 2h,i; Extended Data Fig. 2c,d). Taken together, our data indicate that upon glutamine stress, aPKCs function to relocate Par3 to microtubules, where Par3 has the opportunity to regulate microtubule motor properties, facilitating macropinocytosis.

### Discrete modes of glutamine stress induction differentially regulate the aPKCs

6-Diazo-5-oxo-L-norleucine (DON) is a glutamine analog that broadly inhibits glutamine metabolism and drives metabolic stress that stimulates macropinocytic induction to levels comparable to glutamine deprivation^5^ (Extended Data Fig. 3a-d). As we observed for glutamine depletion, knockdown of the aPKCs or aPKC inhibition via ACPD blocked DON-stimulated macropinocytosis (Extended Data Fig. 3e-l). Also, DON-induced macropinocytosis was abrogated by knockdown of either Par-1a, Par3 or Par6 (Extended Data Fig. 3m-p). Since metabolic stress caused pharmacologically can often mechanistically differ from stress caused by nutrient starvation, we next investigated how these different modes of glutamine stress might impact the regulation of the aPKCs. Interestingly, we found that only DON treatment had the capacity to increase expression of both PKCζ and PKCι proteins and transcripts (Fig. 3a-d). We next examined the phosphorylated forms of PKCζ and PKCι at the T560 and T555 residues, respectively, which are required for the full activation of these kinases ^30–33^. We observed that only glutamine deprivation promoted an increase in the p-aPKC/aPKC ratio (Fig. 3e,f). With DON treatment, although the p-aPKC/aPKC ratio was unaffected, both the phosphorylated and total forms of the proteins were enhanced (Fig. 3e,f). These results suggest that DON modulates PKCζ and PKCι function by increasing their expression, while glutamine starvation primes these kinases for activation.

**Fig. 3.**
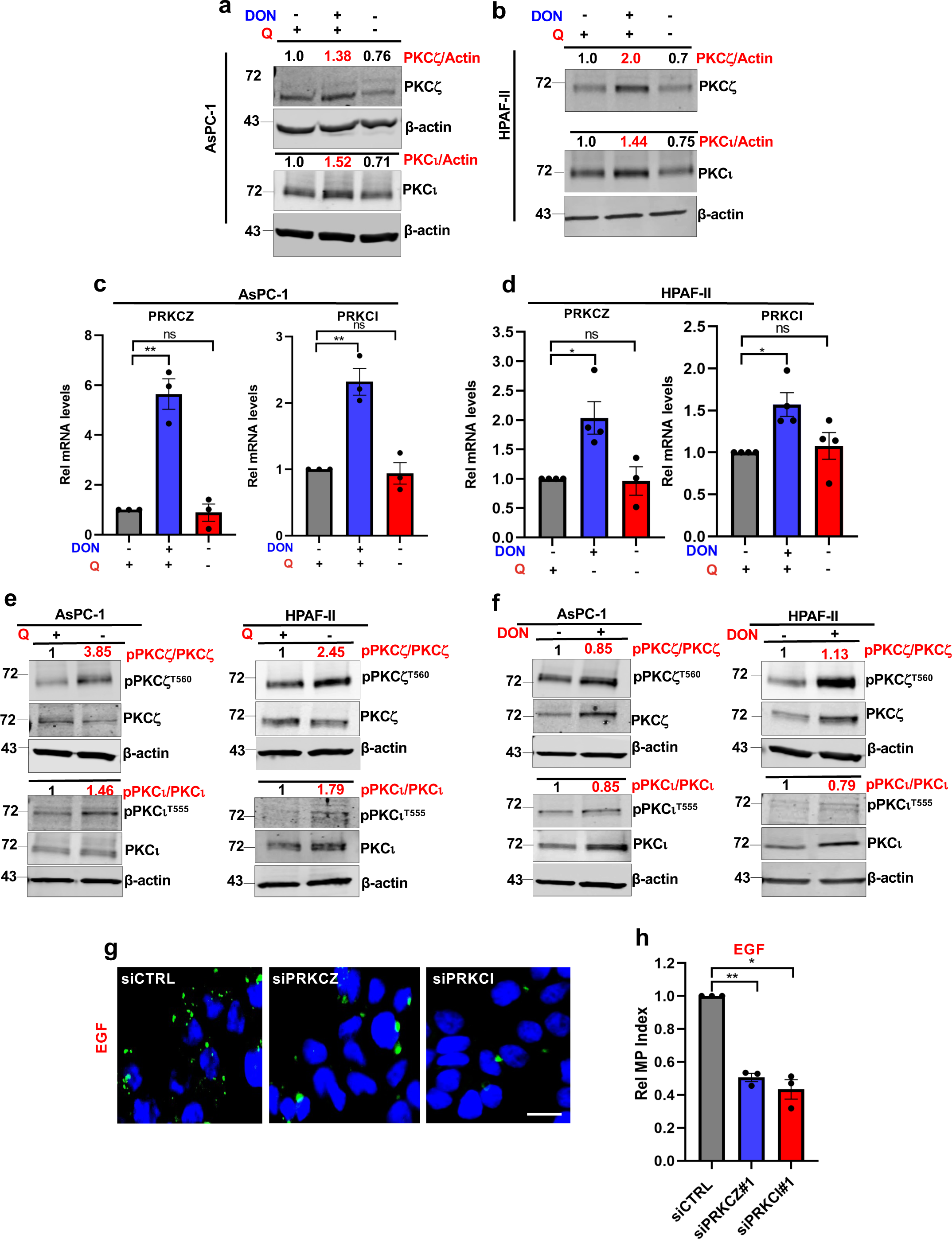
Glutamine deprivation primes aPKCs for activation through the EGFR signaling pathway. **a, b)** Immunoblots assessing PKCζ and PKCι total protein levels in AsPC-1 (a) and HPAF-II (b) cells cultured for 24 hrs in glutamine-replete vehicle (water) control media, 2 mM DON, or in glutamine-free conditions. β-actin was used as a loading control. Data are representative of at least three independent experiments. Densitometry quantifications are presented relative to the vehicle control and normalized to β-actin. **c, d)** Relative *PRKCZ* and *PRKCI* mRNA levels as assessed by RT-qPCR in AsPC-1 (c) and HPAF-II (d) cell cultured in the conditions described in a and b. Data are shown relative to glutamine-replete vehicle control. Data are presented as the mean ± SEM of n=3 or n=4 independent experiments for AsPC-1 or HPAF-II cells, respectively. **e)** Immunoblots assessing levels of the phosphoproteins p-PKCζ^T560^ and p-PKCι^T555^, and total PKCζ and PKCι protein levels in AsPC-1 and HPAF-II cells cultured in glutamine-replete or glutamine-free media. β-actin was used as a loading control. Densitometry quantifications are presented relative to the glutamine-replete condition and values for the phospho-forms are normalized to the total protein. Data are representative of at least three independent experiments. **f)** Immunoblots assessing levels of the phosphoproteins p-PKCζ^T560^ and p-PKCι^T555^, and total PKCζ and PKCι protein levels in AsPC-1 and HPAF-II cells cultured in glutamine-replete media treated with vehicle or 2 mM DON. β-actin was used as a loading control. Densitometry quantifications are presented relative to the glutamine-replete condition and values for the phospho-forms are normalized to the total protein. Data are representative of two independent experiments. **g)** Representative images of macropinocytosis in AsPC-1 cells transfected with the indicated siRNAs and cultured in glutamine-replete media treated with vehicle or EGF at 100 nM for 10 minutes. Scale bar, 20 μm. **h)** Quantification of macropinocytosis for the conditions described in g. Data are shown relative to siCTRL treated with EGF and are presented as mean ± SEM from three independent experiments. Statistical significance was calculated using unpaired two-tailed Student’s t test with Welch’s correction. Ns, non-significant, \**P*<0.05, \*\**P*<0.01.

Glutamine deprivation promotes macropinocytosis through EGFR^5^; therefore, we next explored the interconnectedness between aPKC and EGFR signaling in regulating macropinocytosis upon glutamine starvation. To do this, we took advantage of the fact that stimulation with EGF ligands can mimic glutamine depletion and promote macropinocytosis^5^. Interestingly, we found that knockdown of either PKCζ or PKCι significantly reduced EGF-induced macropinocytosis, suggesting that the aPKCs were indeed functioning to mediate EGFR signaling to drive uptake in PDAC cells (Fig. 3g,h). Depending on the context, there are several different ways that aPKCs can regulate EGFR signaling. In mouse embryonic fibroblasts and lung epithelial cells, upon activation, aPKCs can indirectly influence EGFR signal potentiation by phosphorylating EGFR ligands, priming them for proteolytic cleavage that results in the release of soluble ligand^34, 35^. In glioblastoma cells, aPKC phosphorylation-dependent activation is downstream of EGFR and serves to control glioblastoma progression^36^. In PDAC cells, we found that EGF treatment enhanced aPKC phosphorylation to similar extents as observed with glutamine deprivation (Extended Data Fig. 4a,b). Moreover, aPKC phosphorylation in glutamine-deprived conditions was EGFR-dependent as it was inhibited by erlotinib, an EGFR inhibitor (Extended Data Fig. 4c,d). Interestingly, although DON-induced macropinocytosis is aPKC-dependent, we did not observe enhanced EGFR activation in DON-treated cells, nor was DON-induced macropinocytosis suppressed by erlotinib (Extended Data Fig. 4e-h). Altogether, these data indicate that the mechanistic underpinnings of nutrient stress-driven macropinocytosis might be influenced by how the metabolic stress is achieved, but more importantly, our findings reinforce the importance of the cell polarity proteins in regulating macropinocytosis, independent of the metabolic stress source.

### Glutamine antagonism upregulates aPKC expression through the transcription factor CREM

Having established that priming of the aPKCs is regulated by EGFR signaling to control macropinocytosis in the context of glutamine starvation, we were next interested in examining the mechanistic underpinnings of how aPKC expression is upregulated by DON. Since we observed that DON enhanced the transcript levels of both PKCζ and PKCι, we hypothesized that this increase in expression might be controlled by a specific transcription factor. Using the ChiP Enrichment Analysis (ChEA) dataset, an application that predicts target gene-transcription factor associations by data aggregation from low and high-throughput studies^37^, we generated a list of 29 and 32 transcription factors (TFs) associating with the *PRKCZ* or *PRKCI* promoters, respectively (Extended Data Fig. 5a). For further analyses, we focused on nine TFs that were common to both promoters (Fig. 4a). To determine whether any of these TFs, or any other potential drivers of macropinocytosis, are upregulated by DON, we performed transcriptomics using RNA-Sequencing (RNASeq) in AsPC-1 cells treated with DON or vehicle control. Using this approach, we found that DON upregulated the expression of two of the nine TFs: cyclic-AMP responsive element modulator (CREM) and Rest corepressor 3 (RCOR3; Fig. 4b). As expected, we observed upregulation of *PRKCZ* and *PRKCI* in DON-treated cells, but we also found that other cell polarity transcripts including PAR1A, PARD3 and PARD6B were increased (Extended Data Fig. 5b). This further emphasizes the relevance of these proteins in the glutamine stress response.

**Fig. 4.**
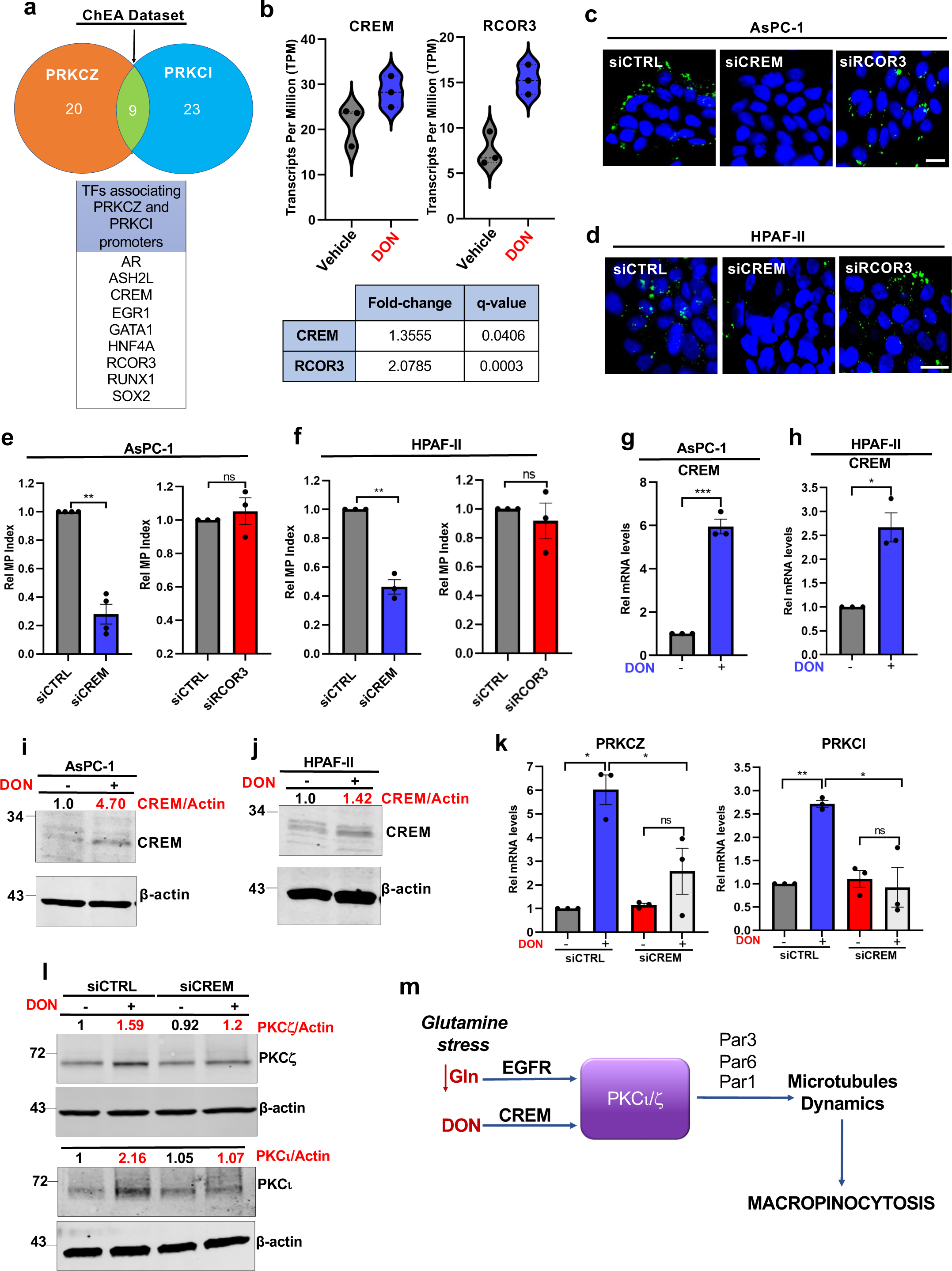
DON upregulates aPKC expression through the transcription factor CREM. **a)** Schematic depicting the transcription factors predicted to bind to the PRKCZ and PRKCI promoters as generated using the ChEA dataset. **b)** Top: Violin plot showing *CREM* and *RCOR3* transcript counts in AsPC-1 cells treated with vehicle or DON (2 mM, 24 hrs) as assessed by RNA-seq. Bottom: Fold-change relative to vehicle and q-values for *CREM* and *RCOR3* transcripts. **c)** Representative images of macropinocytosis in AsPC-1 cells transfected with the indicated siRNAs cultured in glutamine-replete media containing vehicle or 2mM DON. Scale bar, 20 μm. **d)** Representative images of macropinocytosis in HPAF-II cells transfected with the indicated siRNAs cultured in glutamine-replete media containing vehicle or 2mM DON. Scale bar, 20 μm. **e)** Quantification of macropinocytosis for the conditions described in c. Data are shown relative to siCTRL cells treated with DON. Data are presented as the mean ± SEM of n=4 or n=3 independent experiments for siCREM or siRCOR3, respectively. **f)** Quantification of macropinocytosis for the conditions described in d. Data are shown relative to siCTRL cells treated with DON. Data are presented as the mean ± SEM of three independent experiments. **g, h)** Relative *CREM* mRNA levels as assessed by RT-qPCR in AsPC-1 (g) and HPAF-II (h) cultured in glutamine-replete media containing vehicle or DON (2 mM, 24 hrs). Data are presented as the mean ± SEM of three independent experiments. **i, j)** Immunoblots assessing CREM protein levels in AsPC-1 (i) and HPAF-II (j) cells cultured in glutamine-replete media containing vehicle or DON (2 mM, 24 hrs). β-actin was used as a loading control. Data are representative of two independent experiments. Densitometry quantifications are presented relative to vehicle and values are normalized to β-actin. **k)** Relative *PRKCZ* and *PRKCI* mRNA levels as assessed by RT-qPCR in AsPC-1 cells transfected with siCTRL and siCREM and cultured in glutamine-replete media containing vehicle or DON (2 mM, 24 hrs). Data are shown relative to vehicle. Data are presented as the mean ± SEM of three independent experiments. **l)** Immunoblots assessing PKCζ and PKCι total protein levels in AsPC-1 cells cultured in glutamine-replete media containing vehicle or DON. β-actin was used as a loading control. Data are representative of n=3 or n=2 independent experiments for PKCζ or PKCι, respectively. Densitometry quantifications are presented relative to vehicle and values are normalized to β-actin. **m)** Graphical depiction of the mechanistic model outlining how macropinocytosis is controlled by aPKC isoforms in the context of glutamine stress.

To elucidate whether CREM and/or RCOR3 might be involved in DON-stimulated macropinocytosis, we examined uptake in PDAC cells where these TFs were knocked down using specific siRNAs that were validated by RT-qPCR (Extended Data Fig. 5c,d). We observed that depletion of CREM attenuated DON-stimulated macropinocytosis, but RCOR3 did not (Fig. 4c-f). Interestingly, in AsPC-1 cells, macropinocytosis driven by glutamine starvation was unaffected by knockdown of either CREM or RCOR3 (Extended Data Fig. 5e,f). However, in HPAF-II cells, CREM depletion did block macropinocytosis caused by glutamine deprivation, suggesting possible plasticity in the contextual selectivity of CREM in regulating uptake (Extended Data Fig. 5g,h). Consistent with CREM playing a role in DON-mediated responses, we found that CREM transcription and protein expression were enhanced by DON (Fig. 4g-j). We next deciphered whether the observed role of CREM in macropinocytosis is tied to the regulation of aPKC expression. We measured expression of PKCζ and PKCι at the transcript and protein levels in DON-treated conditions in PDAC cells where CREM was knocked down. We determined that CREM depletion suppresses the DON-dependent increases in aPKC transcription (Fig. 4k). Equivalent results were obtained at the protein level (Fig. 4l). These findings indicate that the enhanced aPKC expression that we observed with glutamine stress is modulated by CREM. Moreover, these data suggest that CREM is a novel modulator of the metabolic adaptations, such as macropinocytosis, which occur in response to the pharmacological inhibition of glutamine metabolism. Overall, we found that different modes of glutamine stress seem to activate aPKCs differently, and that multiple cell polarity proteins converge to regulate the microtubule dynamics that control macropinocytosis (Fig. 4m).

### aPKCs support PDAC cell fitness through macropinocytosis

Glutamine is the most depleted amino acid in human PDAC tumors^4, 7^.Therefore, we next assessed the contributions of the aPKCs to PDAC growth and survival under low glutamine conditions. We knocked down either PKCζ or PKCι and analyzed the proliferative capacities of these cells in media with sub physiological levels of glutamine that would be comparable to what is observed in PDAC tumors^4, 5^. In AsPC-1 and HPAF-II cells, we observed that PKCζ or PKCι depletion suppressed proliferation (Fig. 5a,b and Extended Data Fig. 6a,b). It is well established that macropinocytosis functions to support cancer cell fitness through the uptake of extracellular albumin, which serves as a nutrient source to confer a survival advantage to cells when glutamine is limiting^4, 5^. Since we determined that PKCζ and PKCι are integral to macropinocytic stimulation in response to glutamine stress, we hypothesized that aPKC knockdown might suppress the ability of albumin to rescue PDAC cell dependency on free glutamine. Indeed, while albumin supplementation rescued the deleterious effects of glutamine starvation in the control cells, cells with PKCζ or PKCι loss gained no significant advantage to albumin addition (Fig. 5c-f). These data suggested that aPKCs function to sustain PDAC cell survival in glutamine scarcity via macropinocytosis.

**Fig. 5.**
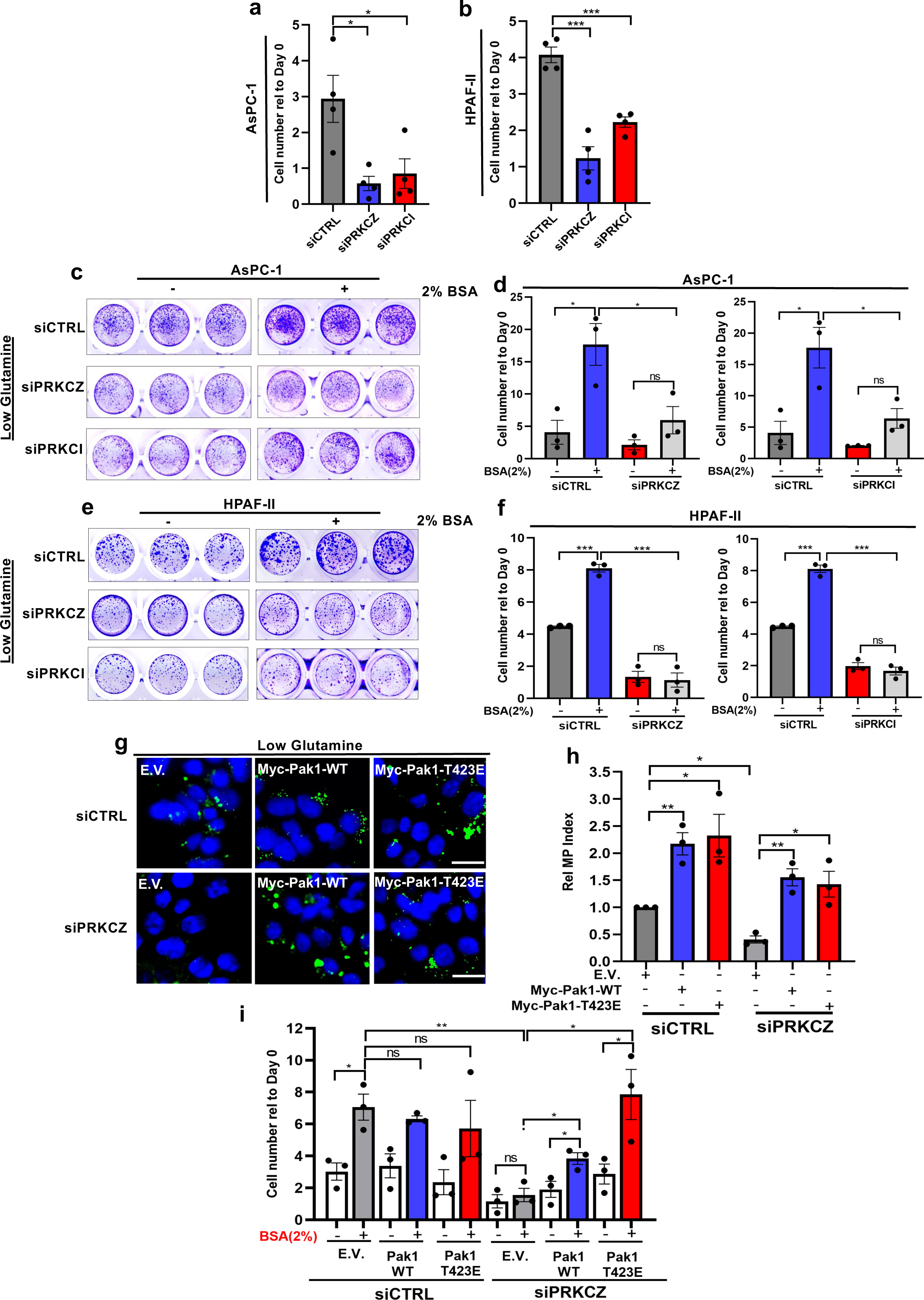
aPKCs modulate PDAC cell fitness and sensitivity to glutamine scarcity through macropinocytosis. **a)** Relative cell growth of AsPC-1 cells transfected with the indicated siRNAs and cultured in low glutamine (0.1 mM) media for 6 days. Cell number was assessed by Syto60 staining and values are presented relative to the day 0 timepoint. Data are presented as the mean ± SEM of four independent experiments. **b)** Relative cell growth of HPAF-II cells transfected with the indicated siRNAs and cultured in low glutamine (0.05 mM) media for 6 days. Cell number was assessed by crystal violet staining and values are presented relative to the day 0 timepoint. Data are presented as the mean ± SEM of four independent experiments. **c)** Representative crystal violet staining for AsPC-1 cells cultured in glutamine-starved conditions with and without supplementation of 2% bovine serum albumin (BSA) for 6 days. **d)** Quantification of cell growth for the conditions described in c. Data are presented relative to the day 0 timepoint. Data are presented as the mean ± SEM of three independent experiments. **e)** Representative crystal violet staining for HPAF-II cells cultured in 0.05 mM glutamine conditions with and without supplementation of 2% BSA for 6 days. **f)** Quantification of cell growth for the conditions described in e. Data are presented relative to the day 0 timepoint. Data are presented as the mean ± SEM of three independent experiments. **g)** Representative images of macropinocytosis in AsPC-1 cells co-transfected with the indicated siRNAs and either empty vector (E.V.), Myc-Pak1-WT or Myc-Pak1-T423E. Cells were cultured in medium containing 0.1 mM glutamine for six days prior to macropinocytosis analysis. **h)** Quantification of macropinocytosis for the conditions described in g. Data are shown relative to siCTRL cells transfected with E.V. Data are presented as the mean ± SEM of three independent experiments. Scale bar, 20 μm. **i)** Quantification of cell growth for the conditions described in g and h with and without supplementation of BSA. Cell number was assessed by crystal violet after 12 days. Data are shown relative to the day 0 timepoint and presented as the mean ± SEM of three independent experiments. Statistical significance was calculated using unpaired two-tailed Student’s t test. Welch’s correction was applied in h. ns, non-significant, \**P*<0.05, \*\**P*<0.01, \*\*\**P*<0.001.

If the proliferative defects observed with aPKC knockdown in low glutamine conditions are attributable to a diminishment in macropinocytic stimulation, then we posited that if macropinocytosis was restored in aPKC knockdown cells, then cell survival would also be restored. Downstream of EGFR, the regulation of nutrient stress-induced macropinocytosis occurs through activation of Pak^5, 38, 39^. In the establishment of apical-basal cell polarity, aPKC function is complemented by Pak1 activity^40^; therefore, we sought to overexpress Pak1 to restore macropinocytosis in aPKC-depleted cells. Control and PKCζ-depleted cells were transfected with an empty vector (E.V.), myc-tagged wild-type Pak1 (Myc-Pak1-WT), or a myc-tagged constitutively active form of Pak1 (Myc-Pak1-T423E). Cells were cultured in low glutamine media, after which macropinocytosis was assessed.

Confirming our previous observations, PKCζ knockdown cells transfected with E.V. displayed lower levels of macropinocytosis compared to control cells (Fig. 5g,h). Importantly, macropinocytosis in the PKCζ-depleted cells was significantly restored upon the expression of either Myc-Pak1-WT or Myc-Pak1-T423E, indicating that Pak1 can compensate for the loss of PKCζ (Fig. 5g,h). Not surprisingly, Pak1 overexpression also enhanced macropinocytosis in the control cells (Fig. 5g,h). PKCζ knockdown and Pak1 overexpression were confirmed by western blot (Extended Data Fig. 6c). After observing the ability of Pak1 to restore macropinocytosis in PKCζ knockdown cells, we asked whether in these conditions the cell viability advantage provided by albumin addition was also restored. To test this, cells were cultured in low glutamine conditions supplemented with or without albumin and cell number was determined by crystal violet staining. In line with our previous results, albumin supplementation did not rescue cell growth in PKCζ knockdown cells (Fig. 5i). We did, however, observe a significant increase in proliferation in PKCζ knockdown cells expressing either of the Pak1 proteins when the media was supplemented with albumin (Fig. 5i). Although macropinocytosis was enhanced in control cells overexpressing Pak1, no additional survival effect was observed. Altogether, these data demonstrate that the function of aPKCs in supporting PDAC cell fitness in the context of nutrient stress is attributable to their role in regulating macropinocytosis.

### aPKCs regulate macropinocytosis *in vivo* and support tumor growth

We next examined whether aPKCs function in tumor growth control and if they regulate macropinocytosis *in vivo.* To knockdown the aPKCs in tumors, we generated stable AsPC-1 cell lines expressing lentiviral-delivered doxycycline-inducible PKCζ or PKCι-targeting shRNAs, as well as a non-targeting shRNA control. We validated that this doxycycline (Dox) system could selectively suppress expression of either PKCζ or PKCι (Extended Data Fig. 7a). We also confirmed that Dox-induced knockdown of either PKCζ or PKCι could inhibit macropinocytosis induced by DON, glutamine starvation or EGF stimulation (Extended Data Fig. 7b-g).

To test aPKC function *in vivo*, we used a heterotopic xenograft mouse model where the stable cell lines were subcutaneously implanted in athymic mice. To induce knockdown, Dox was administered ten days after tumor cell implantation and tumor volumes were calculated based on caliper measurements for approximately five weeks. We observed a significant reduction in tumor growth with either PKCζ or PKCι knockdown relative to controls (Fig. 6a-c). At termination, knockdown in the tumors was confirmed by western blot (Fig. 6d). We observed a decrease in cell proliferation in the PKCζ or PKCι knockdown tumors relative to the controls as assessed by phospho-Histone H3 (p-HistoneH3) immunostaining (Fig. 6e,f). We did not detect a difference in apoptosis within the tumors since levels of cleaved-caspase 3 (Cl-Caspase 3) staining did not change (Extended Data Fig. 8a). To evaluate intratumoral macropinocytosis we used an *ex vivo* macropinocytosis assay that we have previously established^4, 5^. In these tumors, macropinocytosis is low in the periphery and predominantly stimulated within the amino acid-depleted non-peripheral regions of the tumor^5^. In these non-peripheral regions, we found that the macropinocytic index was significantly lower in the PKCζ or PKCι knockdown tumors relative to the control (Fig. 6g,h). Taken together, these data indicate that the aPKCs function in controlling macropinocytosis *in vivo* and are required for tumor growth.

**Fig. 6.**
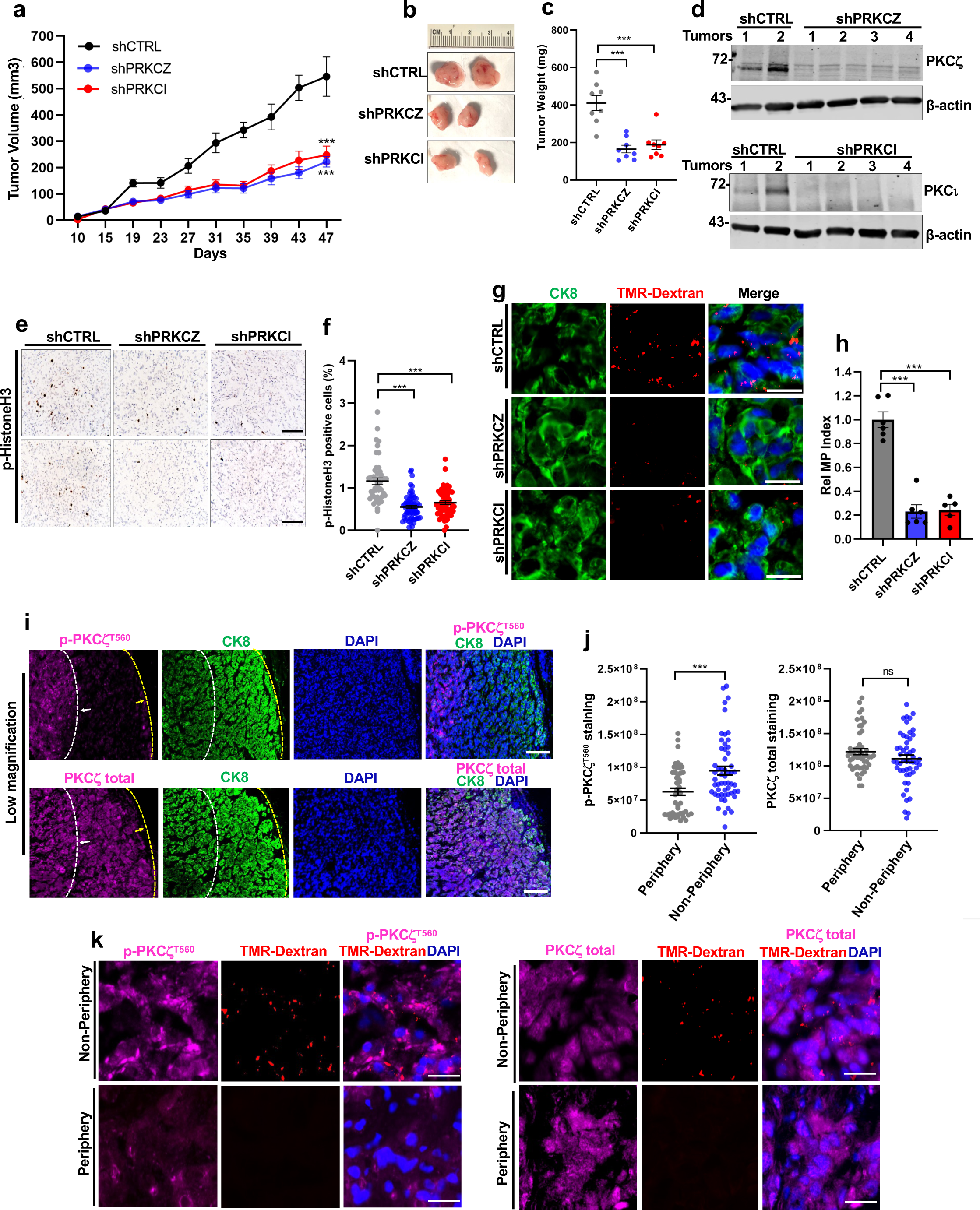
aPKCs support tumor growth and are required for macropinocytosis *in vivo*. **a)** Growth curves for AsPC-1-derived heterotopic xenograft tumors expressing the indicated Dox-inducible shRNAs. Athymic mice were provided Dox in their drinking water and diet. A total of n=8 tumors were measured per condition. **b)** Representative photos of the xenograft tumors at termination. **c)** Measurement of tumor weights at termination for n=8 tumors per condition. **d)** Immunoblots assessing PKCζ and PKCι protein levels in the tumors at termination. Four different knockdown tumors for each aPKC isoform were analyzed relative to two different control tumors. **e)** Representative images of p-HistoneH3 staining in two different tumors per condition. Scale bar, 100 μm. **f)** Quantification of p-HistoneH3 staining in tumor sections. At least 45 image fields were analyzed from n=3 or n=4 tumors for control or aPKC-knockdown conditions, respectively. Data are presented as the mean ± SEM. **g)** Representative images of macropinocytosis from tumors expressing the indicated shRNAs. Macropinosomes are labeled with TMR-dextran (red) and tumor cells were immunostained with CK8 (green). Nuclei are labeled with DAPI (blue). Scale bar, 20 μm. **h)** Quantification of macropinocytosis from the tumors described in g. n=5 tumors were analyzed for each aPKC knockdown group and n=6 tumors were analyzed for the control. Data are shown relative to shCTRL condition. Data are presented the mean ± SEM. **i)** Representative immunofluorescent staining of p-PKCζ^T560^ or total PKCζ (purple) in control tumors. The peripheral and non-peripheral tumor regions were demarcated using the CK8 staining pattern (green). Nuclei are labeled with DAPI (blue). Scale bar, 100 μm. **j)** Quantification of peripheral and non-peripheral staining for p-PKCζ^T560^ and total PKCζ in control tumors. At least 46 fields were analyzed from four different tumors per condition. Data are presented as the mean ± SEM. **k)** Representative images of p-PKCζ^T560^ and total PKCζ (purple) co-staining (purple) with macropinosomes (red) in the peripheral and non-peripheral tumor regions. Nuclei are labeled with DAPI (blue). Scale bar, 20 μm. Statistical significance was calculated using one-way ANOVA with Tukey’s multiple comparison test for a. For c, f, h, and j statistical significance was calculated using unpaired two-tailed Student’s t test. ns, non-significant, \**P*<0.05, \*\**P*<0.01, \*\*\**P*<0.001.

We next sought to explore the link between aPKC activation and nutrient availability *in vivo*. Since glutamine and other amino acids are depleted from the non-peripheral regions of the PDAC tumors^5^, we hypothesized that aPKC phosphorylation might be enhanced in the tumor cores relative to the periphery. Indeed, using immunofluorescence we observed that levels of pPKCζ^T560^ were significantly higher in the non-peripheral regions of the tumors versus the periphery (Fig. 6i,j). This differential pattern was specific to the phosphorylated form since total PKCζ was uniformly expressed throughout the tumors (Fig. 6i,j). This observation was further confirmed by immunohistochemical staining (Extended Data Fig. 8b). Importantly, the tumor cells with enhanced pPKCζ^T560^ levels in the non-peripheral regions of the tumors also stained positive for macropinosomes (Fig. 6k). The antibodies used to detect PKCι proteins did not work well on tissue; therefore, this precluded us from analyzing PKCι in more detail *in vivo*. Altogether, our findings indicate that in the nutrient-depleted regions of the tumor, PKCζ is primed for activation, which in turn enhances the macropinocytic capacity of the cells.

### aPKC expression is upregulated in human PDAC and correlates with poor patient prognosis

Since we observed the difference in the pPKCζ^T560^ levels across the different tumor regions in the AsPC-1 xenografts compared to total PKCζ, we immunostained a tissue microarray containing PDAC specimens from 153 patients that underwent surgical resection. No pPKCζ^T560^ expression was detected in 5 of the specimens, but from the other 148, 125 showed moderate to high expression (Fig. 7a). This observed heterogeneity in pPKC levels in the human tumors is consistent with our observations in the xenografts where pPKC displays intratumoral regional differences. On the other hand, and consistent with our murine data, total PKCζ levels were more homogenous throughout the different specimens (Fig. 7a). To further characterize the cell polarity protein network in PDAC, we interrogated publicly available patient datasets. We found numerous examples where expression of *PRKCI* was upregulated in PDAC relative to normal tissue (Fig. 7b; Extended Data Fig. 9a). Interestingly, we found similar expression correlations with the genes encoding the cell polarity proteins Par6 and Par-1a (Fig. 7c; Extended Data Fig. 9 b,c). In addition, we found that elevated levels of either *PRKCI*, *PRKCZ*, *PARD3* or *PARD6B* were correlated with poor prognosis in PDAC patients (Fig. 7d-g). Altogether, these observations suggest that the aPKC isoforms and/or PAR cell polarity proteins could be potential prognostic indicators in pancreatic cancer.

**Fig. 7.**
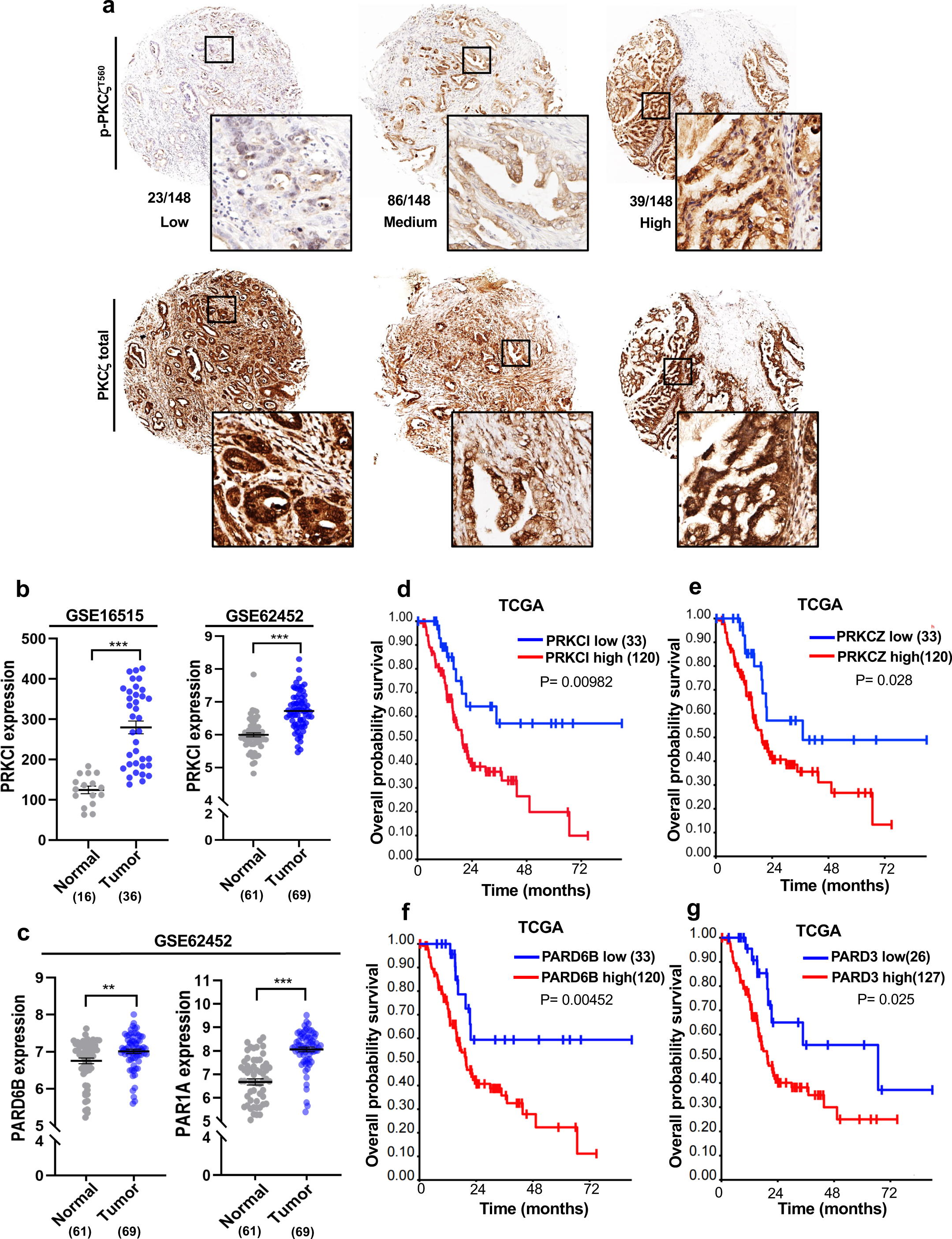
aPKCs and Par polarity proteins are upregulated in PDAC patient tumors, and their high expression correlates with worse prognosis. **a)** Representative immunohistochemical staining for p-PKCζ^T560^ and total PKCζ from clinical samples on a tissue microarray (TMA) containing 152 surgically resected human PDAC specimens. Individual patient samples stained with p-PKCζ^T560^ were pathologically classified according to staining intensity. Images of the TMA samples from the same patient are aligned in columns. **b, c)** *PRKCI (*b) and *PARD6B* and *PAR1A* (c) transcript levels in PDAC human samples and normal pancreatic tissue (adjacent non-neoplastic tissue) from the indicated GSE datasets. Number of patient samples for each group is indicated in parentheses. **d, e)** Overall survival in patients with high and low expression of *PRKCI* (d) and *PRKCZ* (e). Number of patients included in each group is indicated in parentheses. **f, g)** Overall survival in patients with high and low expression of *PARD6B* (f) and *PARD3* (g). Number of patients included in each group is indicated in parentheses. Statistical significance was calculated using unpaired two-tailed Student’s t test for b and c. For d-g, *P* values were determined with the Log-rank test. \**P*<0.05, \*\**P*<0.01, \*\*\**P*<0.001.

## Discussion

The precise functions for the aPKCs in regulating PDAC progression have remained elusive. In this study, we have uncovered a role for the aPKC isoforms PKCζ and PKCι in stimulating macropinocytosis in response to glutamine stress in *KRAS*-mutant PDAC cells, both *in vitro* and *in vivo*. Our findings are consistent with other studies that have described pro-oncogenic roles for these kinases in tumors harboring *KRAS* mutations, such lung adenocarcinoma, basal cell carcinoma, ovarian carcinoma and PDAC^41–48^. Previous studies have not assessed a role for the aPKCs in the metabolic stress response in PDAC, but PKCι has been shown to support PDAC progression through the Rac1-MEK-ERK1/2 signaling axis, and PKCι inhibition leads to impaired tumor growth and a reduction in metastasis and angiogenesis^41^. Interestingly, in a genetically engineered mouse model that is used to study the early stages of tumor initiation, animals of the genotype *LSL-Kras^G12D^; Ptf1a^Cre/+^*, *Prkci^f/f^* still formed pancreatic intraepithelial neoplasia (PanIN) lesions, but these lesions did not progress to invasive adenocarcinoma^46^. Our work demonstrating that PKCι is required for tumor maintenance in established tumors reinforces its attractiveness as a potential therapeutical target in PDAC. For the PKCζ isoform, there is evidence showing that it also participates in Ras-dependent processes, such as the organization of actin cytoskeleton^49^. In the context of *KRAS*-mutant cancers, PKCζ promotes bladder cancer progression inducing the production of reactive oxygen species, that are key mediators of the blebbishield transformation program, an emergency program that allows cancer stem cell resurrection after apoptosis^50^. In PDAC, PKCζ participates in cancer cell motility in association with RhoA GTPase^51^ and favors pancreatic cancer growth and metastasis via activation of STAT3^52^. Thus, PKCζ has also been identified as a positive regulator of PDAC progression. What remains enigmatic is how all these different aspects of cancer biology that are modulated by the aPKCs integrate at the level of tumor progression; however, our observations are in line with these other studies that have established these kinases as pro-oncogenic regulators of tumors harboring *KRAS* mutations. This is in contrast to other oncogenic settings where aPKCs have been postulated to be tumor suppressors^53–58^. Importantly, the precise contribution of each of the isoforms to various aspects of tumor initiation and progression is still poorly understood. Our data suggest that in PDAC, both isoforms independently regulate macropinocytosis since their individual specific reduction show similar effects, but it is likely that in other settings the isoforms might have specific roles since PKCι is known to be widely expressed, whereas PKCζ shows a more restricted expression pattern^59^. One possibility is that the specific roles for each of the isoforms depend on their abundance in the different tissues. For example, during development these kinases are thought to have unique functions since mice deficient in PKCι show early embryonic lethality, whereas no lethality is observed in PKCζ-deficient mice^60,61^. On the other hand, in normal epithelial cells where the aPKCs are both associated with the cell polarity protein network, it has been postulated that the isoforms have similar roles^59, 62^. Future work is required to further elucidate whether there are specific roles or cooperation of the aPKC isoforms in carcinogenesis and cancer progression.

In addition to the aPKCs, we have also identified the cell polarity proteins Par3, Par6 and Par-1a as new regulators of macropinocytosis. Maintenance of cell polarity has been largely associated with tumor suppression mechanisms, such as preventing epithelial-to-mesenchymal transition (EMT)^63^. In this context, aPKCs bind to the scaffold proteins Par3 and Par6 and block EMT through phosphorylation and destabilization of Snail ^64^. In addition, other studies have elucidated that disruption of homeostatic levels of the aPKCs leads to cell polarity loss, which contributes to tumorigenesis^65, 66^. However, acquisition of a cell polarized phenotype is necessary for pro-tumorigenic processes such as cell migration and invasion and this requires the cell polarity protein network to be active^67, 68^. This indicates that in different contexts these proteins can support the progression of cancer. Pro-tumorigenic roles of these proteins have been confirmed in different models. For example, aPKC phosphorylates Par6 to promote EMT and cell migration driven by TGFβ^69^ and Par6 overexpression promotes breast cancer growth^18^. In contrast, Par3 is known to perform dual functions in tumorigenesis, such as in prostate cancer, where it displays pro-tumoral and anti-tumoral effects^70, 71^. In PDAC, loss of Par3 has been associated with increased metastasis, which provided an antitumoral role for Par3, and this antitumoral effect is mediated through the interaction of Par3 with Tiam1^72^, However, the requirement of Par3 that we observed in macropinocytosis and the fact that high Par3 expression associates with poor prognosis in PDAC patients suggest that Par3 can also favor PDAC progression. This suggests that Par3 can display dual functions in pancreatic cancer. Although the loss of these cell polarity proteins leads to the abrogation of macropinocytosis, it is still unclear how in this context this network is regulated. We observed that aPKCs are required for the relocation of Par3 protein to the microtubule network where it is known to regulate microtubule motors, which we show facilitate macropinosome formation. aPKC and Par6 bind through the PB-1 domain of aPKC, forming a complex^59, 69^. It is conceivable that aPKC requires the association with Par6 to modulate macropinocytic uptake and that loss of Par6 might inhibit aPKC function. In cell polarity, aPKCs phosphorylate Par-1a protein at threonine 564 (T564), promoting changes in its subcellular localization^10^. Hence, aPKC might drive subcellular relocation of Par-1a and/or regulate its kinase activity, thereby modifying microtubule dynamics and macropinocytic uptake. Another possibility is that since Par3 is a known target of Par1^73^, Par-1a might be cooperating with aPKCs to modulate Par3 subcellular localization. Future work is required to understand how this cell polarity protein network is coordinated to promote macropinocytosis in the context of nutrient stress. Also, it is important to mention a possible link between aPKC and Pak1 in controlling macropinocytic uptake since both are required for cell polarity establishment^40^, and we observed that Pak1 overexpression compensates for the loss of aPKC in driving macropinocytosis. It is possible that blocking both kinases would lead to a more profound effect on macropinocytosis inhibition, further potentiating the antitumor effects caused by aPKC depletion.

Prior to our work, it remained unexplored whether microtubules have a role in mediating the physical formation of nascent macropinosomes; although, it had been reported that microtubules participate in the post-internalization and trafficking of mature macropinosomes^23^. Here, we have shown that in glutamine stress conditions microtubules are required for macropinocytic uptake, actively participating in the membrane ruffling that drives macropinosome formation. In addition, we observed that inhibition of the motor protein dynein blocks macropinocytic uptake, showing that the retrograde motor function of microtubules is essential in establishing the tension needed to pull the macropinosome inwards, and this might be in part controlled by the accumulation of Par3 in the microtubule network, a process that depends on aPKCs. Microtubules are known to be key elements in the cellular response to metabolic stress^74^. For example, nutrient stress promotes hyperacetylation of tubulin and this turns on the macroautophagy machinery as a compensation mechanism in response to the lack of nutrients^75^. It would not be surprising then that in this context some post-translational modifications promote the rearrangement of the microtubule network to facilitate membrane ruffling, activating the macropinocytic program. Also, microtubules participate in the cellular response to metabolic stress by acting as key sensors of intracellular ATP levels^74, 76, 77^, and it has been shown that low intracellular ATP can be counterbalanced by the macropinocytic uptake of extracellular ATP^78^. Thus, it is possible that low intracellular ATP promotes changes in the microtubule network that would drive the macropinocytic cascade. Future work is essential to further understand how microtubules control macropinocytosis since microtubules would not only be relevant for the formation of macropinosomes, but also in controlling their intracellular trafficking and positioning. A potentially analogous process is how growth factors promote the differential localization of lysosomes in the regulation of mTORC2 and Akt signaling, and how this lysosome positioning is orchestrated by the microtubule network^79^. Hence, it would not be surprising that microtubules control the spatial-temporal positioning of both macropinosomes and lysosomes to permit their fusion and the subsequent digestion and release of the macropinocytic cargo. Uncovering the different post-translational modifications and regulators of the microtubule cytoskeleton that are required for driving macropinocytosis and controlling macropinosome trafficking is essential for a better understanding of how this endocytic pathway is controlled and will shed light on novel effective methods to manipulate its activation and repression.

## METHODS

### Cell culture and reagents

AsPC-1, HPAF-II and HEK-293T cells were obtained from the American Type Culture Collection (ATCC) and were maintained in 100 units/mL penicillin/streptomycin under 5% CO2 at 37°C and routinely tested for mycoplasma contamination using the PCR Mycoplasma Detection Kit (ABM). Cells were cultured in the following media: AsPC-1 in RPMI (Corning) supplemented with 10% fetal bovine serum (FBS), 20 mM HEPES and 1 mM sodium pyruvate and HPAF-II and HEK-293T cells in DMEM (Corning) with 10% FBS and 20mM HEPES. For doxycycline-inducible expression cassettes, Tet System Approved FBS (#631101, Takara) was used. AsPC-1 transduced cells were maintained in 10 μg/mL puromycin (P8833, Sigma).

The following reagents were used: 6-diazo-5-oxo-L-norleucine (DON) (D1242, Sigma); 2-acetyl-1,3-cyclopentanedione (ACPD) (R426911, Sigma); Colchicine (C9754, Sigma); Paclitaxel (T7402, Sigma); Ciliobrevin D (250401, Sigma); Erlotinib (SML2156, Sigma); EGF (E9644, Sigma); Doxycycline (AC446060250, Fisher). For DON treatment, 2mM of DON was added for 24 hours in glutamine-containing serum free media. APCD was used for 3 days at a 4 μM in AsPC-1 cells and 25 μM in HPAF-II cells. Inhibitor was refreshed every 24 hours. Colchicine and paclitaxel were used at 100 ng/mL and 200 nM, respectively, for 18 hours in serum-free media. Ciliobrevin D was used at 50 μM for 30 minutes. For erlotinib experiments, cells were treated for 2 hours at 25 μM. EGF was used at 100 nM at the indicated time-points or for 10 minutes for the macropinocytosis uptake assays. Transduced AsPC-1 cells were treated for 3 days with 2 μg/mL of doxycycline for knock-down induction.

### Glutamine deprivation

Experiments evaluating macropinocytosis in glutamine deprivation were performed in the absence of serum for 24 hours, unless otherwise indicated. Cells were plated in complete culture media, which was exchanged with serum free glutamine-deprived or glutamine-containing media 2–3 days after cell seeding. For glutamine deprivation experiments, glutamine-free RPMI (Corning) or glutamine-free DMEM (Corning) was used. For Figure 5 G and H, cells were cultured in serum-containing media supplemented with 0.1 mM of glutamine for 6 days prior to macropinocytosis assessment.

### siRNA kinome screen

The HT-compatible macropinocytosis assay^13^ was used in conjunction with an siRNA library targeting 679 kinases (siOTP-Dharmacon). Approximately 4,000 AsPC-1 cells were seeded into each siRNA-containing well of 384-well plates and reverse transfected at a final 10nM siRNA concentration. Lipo-RNAimax (ThermoFisher) was used as a transfection reagent. 48 hours after transfection, media was aspirated, and wells were thoroughly washed with sterile PBS (Fisher) and incubated with serum-free glutamine-deprived RPMI media in combination with 2 mM DON for glutamine-stress induction. After 24 hours, macropinocytosis was assessed by a FITC-Dextran (ThermoFisher Scientific) uptake assay. Cells were incubated with media containing 1 mg/mL FITC-Dextran for 30 minutes at 37°C. 4 washes with cold PBS were performed and cells were fixed with 3.7% formaldehyde for 30 minutes. 2 μg/ml of DAPI (MilliPore) was added to stain nuclei. 24 hours later, imaging of the wells was performed in the Sanford Burhnam Prebys Functional Genomics core using a laser-based automated HT microscope (IN Cell 1000, Opera QEHS, Celigo). 2 rounds of screening were performed and macropinocytic index was determined as previously described^13^. Hits showing a reduction greater than 67% were selected for further validation.

### *In vitro* macropinocytosis

Cells were seeded on acid-washed glass coverslips in 24-well plates and subjected to serum starvation with or without nutrient deprivation for 24 hours. For macropinocytosis in siRNA transfected cells, cells were subjected to serum starvation 24 hours after the second transfection, whereas for macropinocytosis in non-transfected or transduced cell lines, serum deprivation was performed 2-3 days after cell plating. After 24 hours of starvation, media was first removed from the wells and the same media containing 1 mg/mL FITC-Dextran (ThermoFisher Scientific) was added back to the cells and plates were incubated for 30 minutes at 37°C. Thereafter, 4-5 washes with cold PBS were done and cells were fixed in 3.7% formaldehyde for 15 minutes. Nuclei were next stained with 2 μg/ml of DAPI(MilliPore) and coverslips were mounted on glass-slides with DAKO-fluorescent media (Agilent Technologies). Images were captured and analyzed using two different approaches. As previously described, pictures were captured at 40X magnification using the EVOS FL Cell Imaging System (Thermo Fisher Scientific) and manually analyzed using ImageJ software (NIH)^5^. The macropinocytic index was calculated by the total particle area per cell. Alternatively, images were automatically captured at 40X magnification using the BioTek Cytation 5 (Agilent Technologies) from the Sanford Burnham Prebys Cell Imaging Core and were subjected to automated analysis using the BioTek Gen5 software^14^. Calculation method of the macropinocytic index was the same as described above. Both approaches have been previously compared and have shown to provide the same results^14^.

### siRNA and plasmid transfection

For macropinocytosis and immunofluorescence assays, cells were transfected in 24-well plates with Lipofectamine RNAimax transfection reagent (ThermoFisher Scientific) and with the corresponding siRNAs at a final concentration of 50 nM following manufacturer’s protocol. For RT-qPCR and Immunoblot analysis, cells were transfected in 12-well and 6-well plates, respectively. Two sequential transfections were performed for each experimental sample. For macropinocytosis and immunofluorescence analysis, serum-free media with the corresponding experimental components was added 24 hours after the second transfection. For knock-down validation, mRNA and protein extracts were collected 48 hours after the second transfection and no serum-starvation was performed. For cell-growth curves, cells were transfected in 6-cm plates and trypsinized 24-hours after second siRNA transfection and seeded in 96-well plates.

The following siRNAs (ThermoFisher Scientific) were used: non-targeting siCTRL (4390844); siPRKCZ#1 (s11128) and #2 (s11129); siPRKCI#1 (s11110) and #2 (s11111); siPAR1-a#1 (s230619) and #2 (s8514); siPARD3 #1 (132893) and #2 (132894); siPARD6B #1 (225936) and #2 (225937); siRCOR3(132395). siCREM (SASI_Hs01_00111489) was purchased from Sigma.

For overexpression assays, cells-were transfected in 6-cm plates 24 hours after second siRNA transfection with Lipofectamine 2000 (ThermoFisher Scientific) and 2 μg of pCMV6M-Pak1-WT (Addgene, 12209), pCMV6M-Pak1-T423E (Addgene, 12208) or empty vector (Origin, pCMV6KN). 24 hours after Pak1 overexpression, cells were trypsinized and seeded in 96-well plates for cell growth analysis.

For lentiviral production, HEK-293T cells were seeded in 6-cm plates and 4.5 ug of Lentiviral vector were cotransfected with 3 ug and 1.5 ug of the packaging vectors psPAX2 and pMD2.G. Lipo2000 (ThermoFisher Scientific) was used as transfection reagent. HEK-293T cells were transfected for 48 hours prior to viral particles collection.

### Immunoblotting

Total protein was extracted from cells or tumor tissues using RIPA buffer (10mM Tris-HCl [pH 8.0], 150mM NaCl, 1% sodium deoxycholate, 0.1% SDS, 1% Triton X-100) with protease and phosphatase inhibitors (Roche). Protein concentrations were measured using the DC Protein Assay Kit (Bio-Rad). 25-30 µg protein samples were run in SDS–PAGE followed by protein transfer using Mini Gel Tank and Mini Blot Module (Life Technologies). Immunoblotting was detected using near-infrared fluorescence (LI-COR) and the Odyssey CLx imager (LI-COR). The following primary antibodies were used: PKCζ (CST 9368S, 1:500), PKCι (BD Transduction Laboratories 610175, 1:700), p-PKCζ^T560^ (Abcam ab62372, 1:2000), p-PKCι^T555-63^ (Abcam ab5813, 1:1000), Par3 (Millipore Sigma 07-330, 1:4000), CREM (Santa Cruz sc-101530, 1:100), Myc (CST 71D10, 1:1000), p-EGFR ^Y1068^ (CST 3777S, 1:1000), EGFR (CST 4267, 1:1000) and β-actin (Sigma A1978, 1:10000). Densitometry analysis was performed with ImageJ software (NIH).

### Quantitative real time PCR (qPCR) and RNA sequencing

Total RNA was extracted from cells with the PureLink RNA Mini Kit (Thermo Fisher Scientific). For quantitative real time PCR (qPCR), cDNA was synthesized using the High-Capacity cDNA Reverse Transcription Kit (Thermo Fisher Scientific). cDNA samples were diluted 1:4 in RNAse free water. qPCR was performed in triplicates with the SYBR Premix Taq II master mix (Takara) and specific primers on the LightCycler 96 (Roche). Relative target gene expression was determined by comparing average threshold cycles (CT) with that of housekeeping genes (*β-actin* or *RPL13*) by the delta-delta CT method. The following primers were used in this study: PRKCZ-F: ATTCATGTTTTCCCGAGCAC; PRKCZ-R: TGTTAAAGCGCTTGGCTTG; PRKCI-F: CGAACAGCTCTTCACCATGA; PRKCI-R: CGTTCTGGTACACAAGGGAAC; PAR1-a F: CAGTCTCCTCAC CACAAAGTGC; PAR1-a-R: TGCTGGTCTGACTCCTTTTCGG; PARD3 F-ATACCAGCTGTCCCCTACAGT; PARD3-R-CTGGACTCAAAGAGCAGTCG; PARD6B-F -GGTGAAGAGCAAGTTTGGAGC; PARD6B-R TAAGCAGTGGATTGGCCGTT; RCOR3-F GTTGGGATGAGAGTCGGAGC; RCOR3-R AACATGCCAAGTGCCTGTTC; CREM-F TGATGGTGTTCAGGGACTGC; CREM-R ATGTCACCAGTGGCAGCTTG; RPL13-F GTTCGGTACCACACGAAGGT; RPL13-R TGGGGAAGAGGATGAGTTTG; β-ACTIN-F ATTGGCAATCAGCGGTTC; β-ACTIN-R CGTGGATGCCACAGGACT.

RNA-Seq was performed by the Sanford Burnham Prebys Genomics Core. AsPC-1 cells were treated with either vehicle or DON for 24 hours in serum-free media. Thereafter, Poly(A) RNA was isolated using the NEBNext Poly(A) mRNA Magnetic Isolation Module (NEB) and barcoded libraries were made using the NEBNext Ultra II Directional RNA Library Prep Kit for Illumina (NEB). Libraries were pooled and single-end sequenced (1×75 bp) on the Illumina NextSeq 500 using the High-Output V2 Kit (Illumina). Read data were processed in BaseSpace, and reads were aligned to Homo sapiens genome (hg19) using STAR aligner with default settings.

### Immunofluorescence

Cells seeded on acid-wash coverslips were washed with cold PBS and fixed with 4% Paraformaldehyde (PFA) for 15 minutes at room temperature. Cells were then permeabilized with 0.2%-Triton-X-100 in PBS for 10 minutes and blocked in 10% Normal Goat Serum in PBS for 1 hour. Thereafter, cells were stained overnight with primary antibodies for tubulin (Sigma T6074, 1:500) and Par3 (Sigma HPA030443, 1:50) diluted in blocking buffer. Three washes of PBS were performed, and cells were incubated with secondary antibodies Alexa Fluor 488 (ThermoFisher Scientific A32723, 1:1000) and Alexa Fluor 594 (ThermoFisher Scientific A11012, 1:1000) for tubulin and Par3 staining, respectively. Secondary antibody solutions were supplemented with 2μg/mL of DAPI (Millipore Sigma) for nuclear staining. Finally, cells were washed three times with PBS and mounted on glass - slides using fluorescent-DAKO mounting media (Agilent Technologies). Images of tubulin morphology and costainings of tubulin and Par3 were captured at the Sanford Burnham Prebys Cell Imaging Core, using a Zeiss LSM 710 laser scanning confocal microscope equipped with a Plan Apochromat 63X/1.4 NA oil immersion objective. Quantification of the Par3 puncta/cytosolic fraction ratios was performed with the ImageJ software (NIH). Cytosolic fraction was calculated subtracting the intensity (mean gray value*total area) of the Par3 puncta to the total Par3 intensity (Cytosol _intensity_=Total Par3_intensity_-Puncta Par3_intensity)._ Percentage of Par3 puncta/cytosol was determined dividing Par3 puncta intensity to the intensity of the cytosolic fraction ((Puncta Par3 _intensity_/ Cytosol _intensity_) *100).

Tumor sections frozen in OCT compound and injected with TMR-dextran were first sectioned by the SBP Histology Core. Samples were then thawed at room temperature for 5-10 minutes and fixed with 3.7% formaldehyde for 15 minutes. Slides were permeabilized with 0.2%-Triton-X-100 in PBS for 10 minutes and blocked in 10% Normal Goat Serum and 1% BSA in PBS for 1 hour. After three washes with PBS, samples were incubated with primary antibodies for p-PKCζ^T560^ (Abcam ab62372, 1:400); PKCζ (Abcam ab59364, 1:400); CK8 (TROMO-I, DSHB, 1:400). After three PBS washes, slides were incubated with the Alexa-Fluor 488 (ThermoFisher Scientific A11006, 1:1000) and Alexa-Fluor 647 (ThermoFisher Scientific, A32733, 1:1000) for CK8 and p-PKCζ/ PKCζ staining, respectively. Secondary antibody solutions were supplemented with 2 μg/mL of DAPI (Millipore Sigma). Pictures were captured using an EVOS M5000 Cell Imaging System (ThermoFisher Scientific) equipped with a Plan Fluorite 40x/0.75 NA air objective. Intensity of p-PKCζ and total PKCζ staining in the peripheral and non-peripheral regions of the tumor was quantified with ImageJ software (NIH). Tumor periphery was established by the presence of non-tumor cells (CK8 negative) at the edges of the tumors. For single CK8 immuno-staining, no permeabilization was performed and primary antibody was incubated for 4 hours at the same dilution.

### Growth curves

After 24 hours of the second siRNA transfection or Pak1 overexpression, cells were trypsinized and 5,000 cells were seeded in wells of 96-well plates. 24 hours after cell plating, time 0 of the growth curve was determined and media was replaced with serum-containing media with 0 or 0.1 mM glutamine for AsPC-1 cells and 0 or 0.05 mM glutamine for HPAF-II cells, supplemented and non-supplemented with 2% serum-albumin (126609, Millipore). Media was refreshed every 3 days for cell growth analysis in knock-down cells, and every 5-6 days for experiments of PKCζ knock-down with Pak1 overexpression. Cell growth curves in AsPC-1 cells were established at 0.1 mM glutamine conditions and at 0.05 mM glutamine for HPAF-II cells. For BSA supplementation experiments in aPKC knock-down cells, BSA rescues were determined in AsPC-1 cells cultured in 0 mM glutamine and in 0.05 mM glutamine for HPAF-II cells. BSA rescues in AsPC-1 PKCζ knock-down cells with Pak1 overexpression were determined at 0.1 mM glutamine cell culture conditions. Cell growth in AsPC −1 and HPAF-II aPKC knock-down cells was determined at day 3 and day 6 after time 0 establishment. For BSA rescue experiments in aPKC knock-down cells, cell growth was determined at day 6 and for BSA rescues in AsPC-1 knock-down cells overexpressed with Pak1, growth was determined at day 12.

For cell growth analysis with crystal violet, media was aspirated, and cells were stained with 0.5% crystal violet for 30 minutes and thoroughly washed in tap-water. When plates were dry, these were scanned, and relative number was calculated by determining the stained area with ImageJ software (NIH). For Syto60 staining, wells were aspirated and incubated for 45 minutes with 1 μM Syto60 (S11342, ThermoFisher) in 0.2% Triton X-100 in PBS. Wells were washed in PBS and plates were imaged at 700 nm on the Odyssey CLx imager (LI-COR). Relative cell number was calculated as described above. Three replicates per condition were used in each independent experiment.

### Lentiviral production and transduction

Tet-pLKO-puro (Addgene plasmid #21915) constructs for doxycycline-induced shRNA expression were generated according to the manufacturer’s instructions. For constructs generation, the following oligonucleotides sequences were used: sh*PRKCZ*#1 (TRCN0000001221); sh*PRKCZ#2* (TRCN0000199469); sh*PRKCI*#1 (TRCN0000006039); sh*PRKCI*#2 (TRCN0000006041) from the TRC database. Sequences and cloning procedure for the non-targeting shRNA were previously described^80^.

Lentiviral production was carried out as described above. 48 hours after transfection, lentiviral particles were collected, and medium containing viral particles was filtered with a 0.45 μm filter (Fisher). Cells were transduced with virus-containing media and 8μg/ml Polybrene (MilliporeSigma). 48 hours after transduction, media containing 10 μg/ml puromycin were added for selection. Infected cells were maintained in media supplemented with 10 μg/ml puromycin for further use.

### Animal studies

Female nude mice (Foxn1^nu^/Foxn1^nu^) aged between 6 to 7 weeks old were purchased from The Jackson Laboratories. All animals were housed in sterile caging and maintained under pathogen-free conditions. The Sanford Burnham Prebys Medical Discovery Institute (SBP) IACUC policy limits tumors in mice to 2 cm^3^ and this was not exceeded. Mice were housed four per cage in individually ventilated cage systems. Relative humidity in the rodent rooms is monitored and recorded daily by animal technicians but is not controlled. Generally, animal room RH (relative humidity) is within the acceptable range of 30-70% for rodents. A standard diurnal light cycle of 12-hr light: 12-hr dark was used with a rodent room thermostat temperature set point of 72°F.

One million cells were resuspended in a 1:1 dilution of PBS and Matrigel (Corning) and subcutaneously injected in both flanks of the mice. Tumor growth was monitored using an electronic caliper and tumor volume was determined according to the *V* = (*L* × *W*^2^)/2 equation. 10 days after cell inoculation, doxycycline was supplemented in the diet (TD.01306, Envigo) and in the drinking water containing 2 mg/mL of doxycycline and 5% sucrose. Drinking water and diet were changed twice and once a week, respectively. Mice were euthanized 27-30 days after tumor cells injection and tumors were sliced and accordingly processed for immunohistochemistry, immunofluorescence and macropinocytosis analysis. A portion of tissue was snap-frozen in liquid nitrogen for immunoblot analysis.

### *Ex vivo* macropinocytosis

As previously described^81, 82^, sections with ∼5 mm thickness were cut from fresh tumor tissues and injected at different sites with 20 mg/mL 10-kDa TMR-Dextran (ThermoFisher Scientific) and immersed in 400-500 μL TMR-Dextran for 15 minutes. After quickly washing three times with PBS, cross sections were embedded in OCT compound and snap-frozen on dry ice. Frozen sections were cut by the SBP Histology Core and subjected to CK8 immunofluorescence staining as previously explained. Images were captured at 40X magnification using the EVOS FL Cell Imaging System (ThermoFisher Scientific) and analyzed with the ImageJ software (NIH) as previously described^5^.

### Immunohistochemistry

Xenografts were fixed in 10% formalin. Fixed tissue was embedded in paraffin and sectioned by the SBP Histology Core. Antigen retrieval was performed by microwave-heating in 10 mM sodium citrate (pH 6) and endogenous peroxidases were quenched in 3% hydrogen peroxide. After that, sections were permeabilized for 10 minutes with 0.2%-Triton X-100 in TBS containing 0.1% Tween-20 (TBS-T), blocked in 2% BSA, 10% goat serum in TBS-T for 1 hour at room temperature and then incubated with primary antibodies diluted in 2% BSA/TBS-T overnight at 4°C. After washes, sections were incubated with biotinylated goat anti-rabbit secondary antibody (1:1000, Vector, BA-1000-1.5) for 1.5 hours at room temperature followed by incubation with the VECTASTAIN Elite ABC HRP Kit (Vector Labs) and the DAB HRP Substrate Kit (Vector Labs). Nuclear counterstaining was performed by hematoxylin staining. Images were captured with a brightfield Olympus CX-31 microscope coupled with INFINITY camera and INFINITY capture software (Lumenera). The following primary antibodies and dilutions were used: p-Histone H3 (CST, 9701,1:200,); p-PKCζ^T560^ (Abcam ab62372, 1:400); PKCζ (Abcam ab59364, 1:400) and cleaved caspase 3 (CST 9664,1: 1,000) For the pHis-H3, the number of p-HistoneH3-positive nuclei per image field was determined using the Fiji software (NIH).

### Clinical data analysis

Immunohistochemical staining surgically resected PDAC (152 patients) was performed using p-PKCζ^T560^ (Abcam ab62372, 1:400) and PKCζ (Abcam ab59364, 1:400) antibodies on a previously described PDAC TMA (quote PMID:20142597) that was provided by the Hirshberg Foundation funded UCLA Pancreas Bank. p-PKCζ^T560^ staining was independently scored by a gastrointestinal pathologist (DWD) based on cytoplasmic intensity (0-3; 0-absent; 1-low; 2-moderate; 3-high) on multiple (2–3) cores for each tumor. The final IHC score for each tumor was calculated as the average intensity across its multiple cores. PKCζ staining intensity was uniformly high (3+) across all tumor cores and therefore not individually scored or evaluated.

PDAC patient data for expression analysis were obtained from the following GSE datasets: GSE62452, GSE28735, GSE15471, GSE16515, [GSE183795](https://www.ncbi.nlm.nih.gov/geo/query/acc.cgi?acc=GSE16515). To analyze survival correlations, patient data was accessed through the R2: Genomics Analysis and Visualization Platform (http://r2.amc.nl) and the “Mixed pancreatic adenocarcinoma (2022vs32)” dataset from TCGA was selected. Overall survival correlation with PRKCI, PRKCZ, PARD3 and PARD6B expression levels was assessed using the following setting: cut_off modus = scan, minimal group size = 8. Cut off was established based on significance.

### Quantification and statistical analysis

All graphs were made using GraphPad Prism software (GraphPad). Results are shown as the mean of at least three independent experiments ± standard error of the mean (SEM) unless stated otherwise. Statistical significance was determined by the unpaired two-tailed Student’s t test with Welch’s correction when appropriate, except for tumor growth analysis, that was determined by one-way ANOVA followed by Tukey test for multiple comparisons. P values less than 0.05 were considered statistically significant (*p < 0.05, **p < 0.01, ***p < 0.001 and ns=not significant).

## ACKNOWLEDGEMENTS

We thank all members of the Commisso laboratory for their helpful comments, discussions, and scientific advice. This work was supported by NIH grants R01CA254806 and R01CA207189 to C.C. Sanford Burnham Prebys Medical Discovery Institute core services are supported by NCI Cancer Center Support grant P30CA030199.

## AUTHOR CONTRIBUTIONS STATEMENT

G.L. and C.C. designed the study and wrote the manuscript. G.L. performed most of the experimental work and prepared all the figures. S-W.L. and P.A.-B. performed the siRNA kinome-wide screen. S-W.L. performed validation for the hits from the siRNA screen. K.D.-P. performed the immunohistochemical staining of the TMA. S.M. assisted with the immunohistochemical staining of the tumor xenograft sections. D.W.D provided the TMA samples and scored the stained TMAs. C.C. supervised the study and obtained funding. All authors reviewed, edited, or commented on the manuscript.

## COMPETING INTESTEST STATEMENT

C.C. is an inventor on a U.S. patent titled ‘‘Cancer diagnostics, therapeutics, and drug discovery associated with macropinocytosis,’’ Patent number: US-11209420-B2.

---

**Extended Data Fig. 1.**
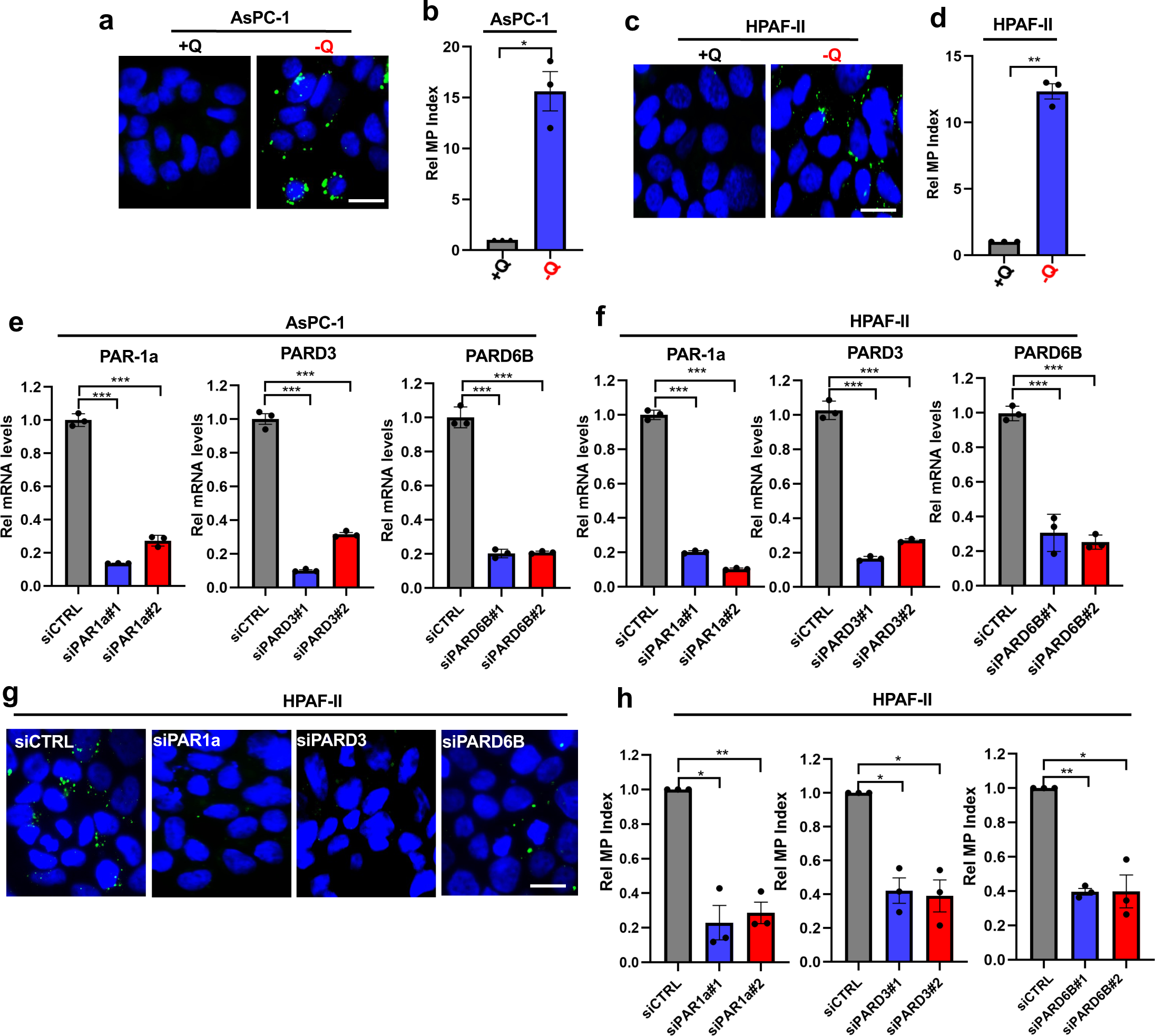
Glutamine starvation stimulates macropinocytosis through cell polarity proteins. **a)** Representative images of macropinocytosis in AsPC-1 cells cultured in glutamine-replete (+Q) or glutamine-starved (-Q) media. Scale bar, 20 μm. **b)** Quantification of macropinocytosis for the conditions described in a. Data are shown relative to the +Q condition and are presented as the mean ± SEM of three independent experiments. **c)** Representative images of macropinocytosis in HPAF-II cells cultured in glutamine-replete (+Q) or glutamine-starved (-Q) medium. Scale bar, 20 μm. **d)** Quantification of macropinocytosis for the conditions described in c. Data are shown relative to the +Q condition and are presented as the mean ± SEM of three independent experiments. **e, f)** Relative *PAR1A, PARD6B* and *PARD3* transcripts levels in AsPC-1 (e) and HPAF-II (f) cells transfected with non-targeting siCTRL or siRNAs targeting *PAR1A*, *PARD6B* or *PARD3*. Data are shown relative to siCTRL. Data are presented as the mean ± SEM and are representative of at least three independent experiments. **g)** Representative images of macropinocytosis in glutamine-starved HPAF-II cells transfected with siCTRL or siRNAs targeting *PAR1A*, *PARD6B* or *PARD3*. Scale bar, 20 μm. **h)** Quantification of macropinocytosis in HPAF-II cells transfected with siCTRL or siRNAs targeting *PAR1A*, *PARD6B* or *PARD3*. Data are shown relative to siCTRL cell cultured in glutamine-free conditions and are presented as the mean ± SEM of three independent experiments. Statistical significance was calculated using unpaired two-tailed Student’s t test with Welch’s correction. \**P*<0.05, \*\**P*<0.01, \*\*\**P*<0.001.

**Extended Data Fig. 2.**
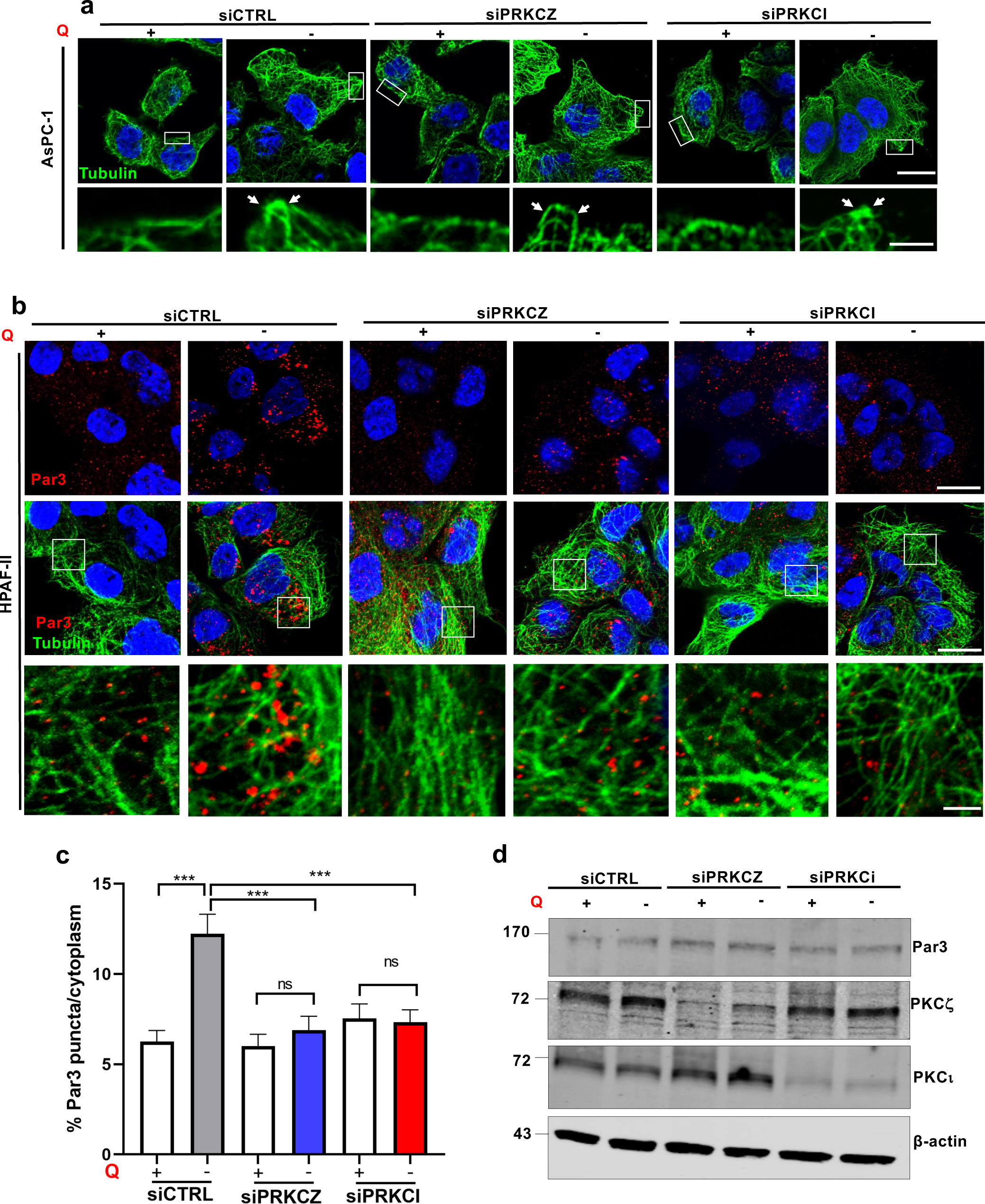
Glutamine stress reorganizes microtubules independently of the aPKCs. **a)** Representative images of immunofluorescence staining of microtubules (green) in AsPC-1 cells transfected with siCTRL, siPRKCZ#1 or siPRKCI#1. Cells were cultured in glutamine-replete (+Q) or glutamine-free (-Q) media. Nuclei are labeled with DAPI (blue). Scale bar, 20 μm. Bottom row are higher magnification images of the boxed areas. Scale bar 5 μm. White arrows indicate the membrane ruffles. Images are representative of at least three independent experiments. **b)** Representative immunofluorescent images of Par3 localization (red) relative to the microtubule network (green) in HPAF-II cells transfected with siCTRL, siPRKCZ#1 or siPRKCI#1. Cells were cultured in glutamine-replete or glutamine-free media. Nuclei are labeled with DAPI (blue). Scale 20 μm. Bottom row are higher magnification images of the boxed areas. Scale bar, 5 μm. **c)** Quantification of the percent of Par3 protein found in subcellular puncta versus the cytoplasm. At least n=50 cells per condition were analyzed. Data are presented as mean ± SEM from three independent experiments. **d)** Immunoblot of Par3, PKCζ and PKCι proteins in lysates from cells cultured in the conditions described in b. β-actin was used as a loading control. Data is representative of three independent experiments. Statistical significance was calculated using unpaired two-tailed Student’s t test. ns, non-significant, \*\*\**P*<0.001.

**Extended Data Fig. 3.**
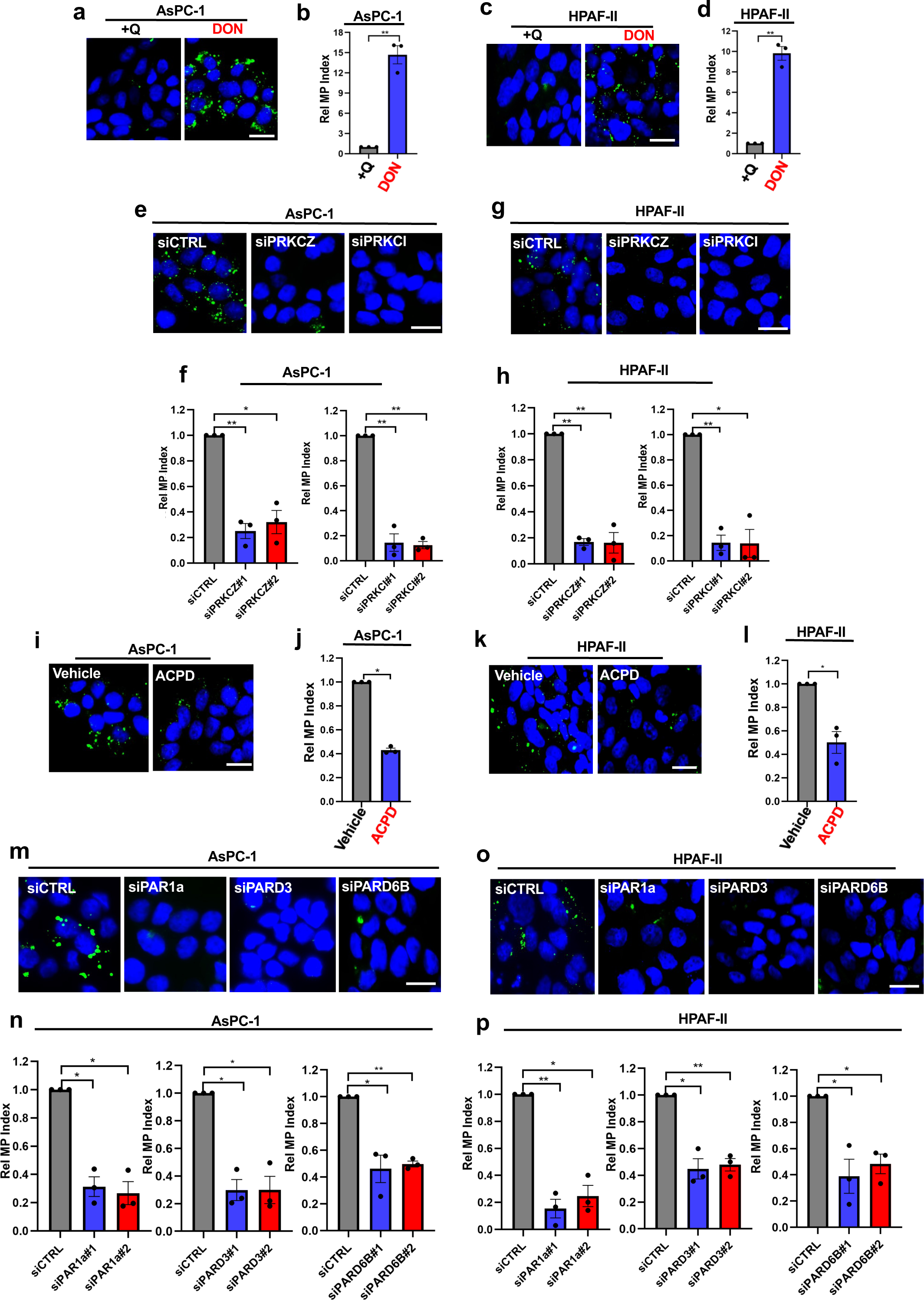
DON induces macropinocytosis through cell polarity proteins. **a)** Representative images of macropinocytosis in AsPC-1 cells cultured in glutamine-replete medium treated with vehicle (water) or DON (2 mM, 24 hrs). Scale bar, 20 μm. **b)** Quantification of macropinocytosis for the conditions described in a. Data are shown relative to the vehicle control and are presented as the mean ± SEM of three independent experiments. **c)** Representative images of macropinocytosis in HPAF-II cells cultured in glutamine-replete medium treated with vehicle or DON. Scale bar, 20 μm. **d)** Quantification of macropinocytosis for the conditions described in c. Data are shown relative to the vehicle control and are presented as the mean ± SEM of three independent experiments. **e)** Representative images of macropinocytosis in DON-treated AsPC-1 cells transfected with siCTRL, siPRKCZ or siPRKCI. Scale bar, 20 μm. **f)** Quantification of macropinocytosis for the conditions described in e. Data are shown relative to siCTRL and are presented as the mean ± SEM of three independent experiments. **g)** Representative images of macropinocytosis in DON treated HPAF-II cells transfected with siCTRL, siPRKCZ or siPRKCI. Scale bar, 20 μm. **h)** Quantification of macropinocytosis for the conditions described in g. Data are shown relative to siCTRL and are presented as the mean ± SEM of three independent experiments. **i)** Representative images of macropinocytosis in DON-treated AsPC-1 cells treated with vehicle or ACPD at 4 μM for 72 hrs. Scale bar, 20 μm. **j)** Quantification of macropinocytosis for the conditions described in i. Data are shown relative to vehicle control and are presented as the mean ± SEM of three independent experiments. **k)** Representative images of macropinocytosis in DON-treated HPAF-II cells treated with vehicle or ACPD at 25 μM for 72 hrs. Scale bar, 20 μm. **l)** Quantification of macropinocytosis for the conditions described in k. Data are shown relative to vehicle control and are presented as the mean ± SEM of three independent experiments. **m)** Representative images of macropinocytosis in DON-treated AsPC-1 cells transfected with siCTRL or siRNAs targeting *PAR1A*, *PARD3* or *PARD6B*. Scale bar, 20 μm. **n)** Quantification of macropinocytosis for the conditions described in m. Data are shown relative to siCTRL and are presented as the mean ± SEM of three independent experiments. **o)** Representative images of macropinocytosis in DON-treated HPAF-II cells transfected with siCTRL or siRNAs targeting *PAR1A*, *PARD3* or *PARD6B*. Scale bar, 20 μm. **p)** Quantification of macropinocytosis for the conditions described in o. Data are shown relative to siCTRL and are presented as the mean ± SEM of three independent experiments. Statistical significance was calculated using unpaired two-tailed Student’s t test with Welch’s correction. \**P*<0.05, \*\**P*<0.01.

**Extended Data Fig. 4.**
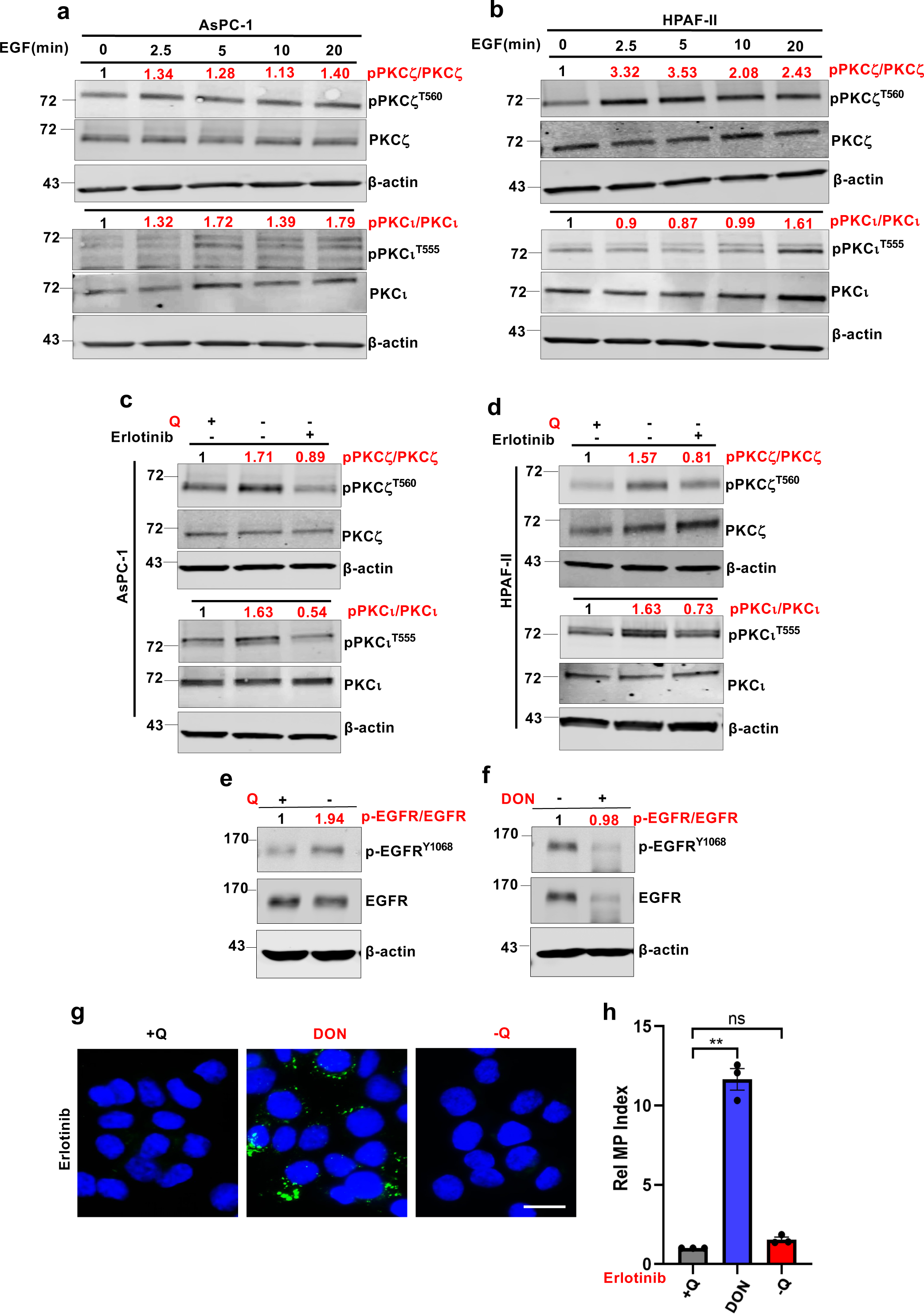
EGFR signaling pathway primes aPKC isoforms for activation in glutamine deprivation, but it is not involved in macropinocytosis promoted by DON. **a, b)** Immunoblots assessing levels of p-PKCζ^T560^, p-PKCι^T555^ and total PKCζ and PKCι protein in AsPC-1 (a) and HPAF-II cells (b) treated with EGF (100 nM) for the indicated times. β-actin was used as a loading control. Densitometry quantifications are presented relative to the time 0 timepoint and values for the phospho-forms are normalized to the total protein. Data are representative of at least three independent experiments. **c, d)** Immunoblots assessing levels of p-PKCζ^T560^, p-PKCι^T555^ and total PKCζ and PKCι protein in AsPC-1 (c) and HPAF-II cells (d) cultured in glutamine-replete or glutamine-free media treated with vehicle or erlotinib (25 μM, 2 hrs). β-actin was used as a loading control. Densitometry quantifications are presented relative to the glutamine-replete condition and values for each phospho-forms are normalized to the total protein. Data are representative of two independent experiments. **e)** Immunoblots assessing levels of p-EGFR^Y1068^ and total EGFR protein in AsPC-1 cells cultured in glutamine-replete and glutamine-free media. β-actin was used as a loading control. Densitometry quantifications are presented relative to the glutamine-replete condition. **f)** Immunoblots assessing levels of p-EGFR^Y1068^ and total EGFR protein in AsPC-1 cultured in glutamine-replete media and treated with vehicle or DON. β-actin was used as a loading control. Densitometry quantifications are presented relative to the vehicle control and values for each phospho-form are normalized to the total protein. **g)** Representative images of macropinocytosis in erlotinib-treated AsPC-1 cells cultured in glutamine-replete media treated with vehicle or DON and in cells cultured in glutamine-free media. Scale bar, 20 μm. **h)** Quantification of macropinocytosis for the conditions described in g. Data are shown relative to the vehicle control and are presented as the mean ± SEM of three independent experiments. Statistical significance was calculated using unpaired two-tailed Student’s t test with Welch’s correction. ns, non-significant, \*\**P*<0.01.

**Extended Data Fig. 5.**
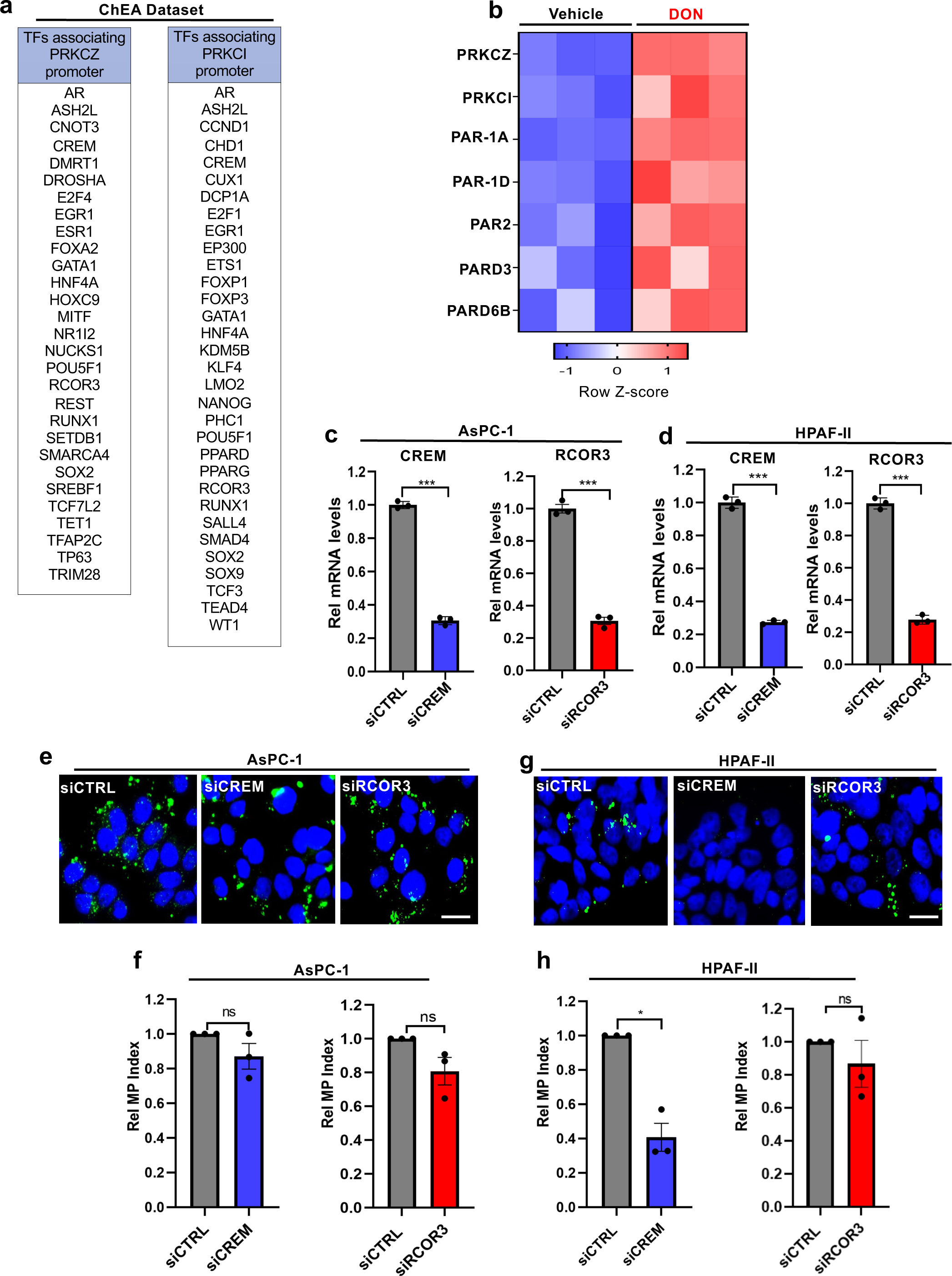
Cell polarity-related transcripts are upregulated in DON-treated cells. **a)** List of transcription factors predicted to bind to the *PRKCZ* and *PRKCI* promoters as generated using the ChEA dataset. **b)** Heat-map of differential gene expression for genes involved in cell polarity regulation in vehicle and DON-treated AsPC-1 cells (in triplicate) as generated by our RNA-seq dataset. **c, d)** Relative *CREM* and *RCOR3* transcript levels in AsPC-1 (c) and HPAF-II (d) cells transfected with siCTRL, siCREM or siRCOR3. Data are shown relative to siCTRL. Data are presented as the mean ± SEM and are representative of at least three independent experiments. **e)** Representative images of macropinocytosis in glutamine-starved AsPC-1 cells transfected with the indicated siRNAs. Scale bar, 20 μm. **f)** Quantification of macropinocytosis for the conditions described in e. Data are shown relative to siCTRL and are presented as the mean ± SEM of three independent experiments. **g)** Representative images of macropinocytosis in glutamine-starved HPAF-II cells transfected with the indicated siRNAs. Scale bar, 20 μm. **h)** Quantification of macropinocytosis for the conditions described in g. Data are shown relative to siCTRL and are presented as the mean ± SEM of three independent experiments. Statistical significance was calculated using unpaired two-tailed Student’s t test with Welch’s correction. ns, non-significant, \**P*<0.05, \*\*\**P*<0.001.

**Extended Data Fig 6.**
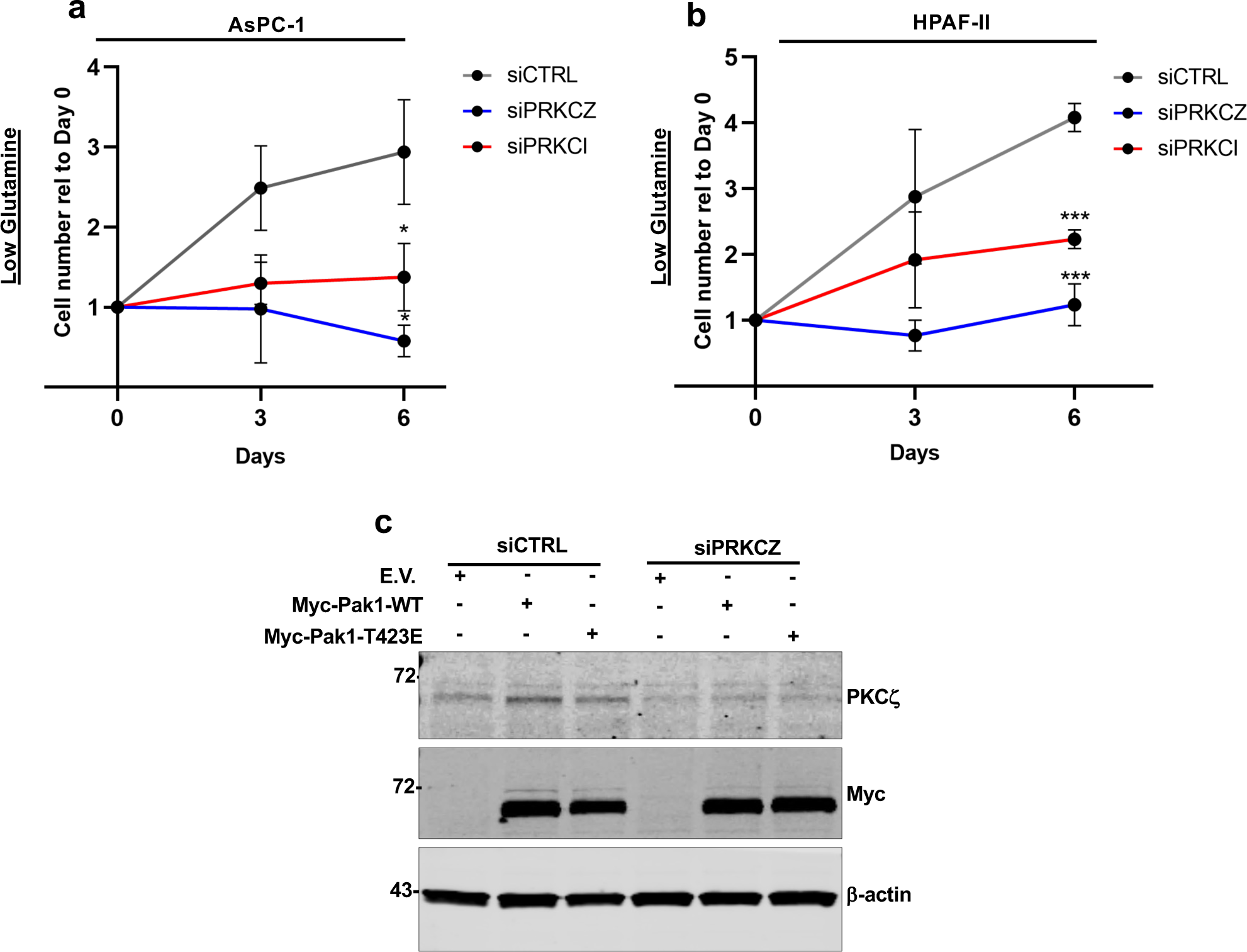
aPKCs regulate cell proliferation in low glutamine conditions. **a)** Relative cell growth of AsPC-1 cells transfected with siCTRL, siPRKCZ or siPRKCI and cultured in low-glutamine (0.1 mM) for 6 days. Cell number was assessed by Syto60 staining on days 0, 3 and 6. Data at each timepoint are shown relative to day 0. Data are presented as the mean ± SEM of four independent experiments. **b)** Relative cell growth of HPAF-II cells transfected with siCTRL, siPRKCZ or PRKCI and cultured in low-glutamine (0.05 mM) for 6 days. Cell number was assessed by crystal violet staining on days 0, 3 and 6. Data at each timepoint are shown relative to day 0. Data are presented as the mean ± SEM of four independent experiments. **c)** Immunoblots assessing PKCζ knock-down in AsPC-1 cells expressing siCTRL or siPRKCZ#1. Cells were also transfected with either empty vector (E.V.), Myc-Pak1-WT or Myc-Pak1-T423E and cultured for six days prior to analysis. β-actin was used as a loading control. Statistical significance was calculated using unpaired two-tailed Student’s t test at day 6 of the growth curve. \**P*<0.05, \*\*\**P*<0.001.

**Extended Data Fig. 7.**
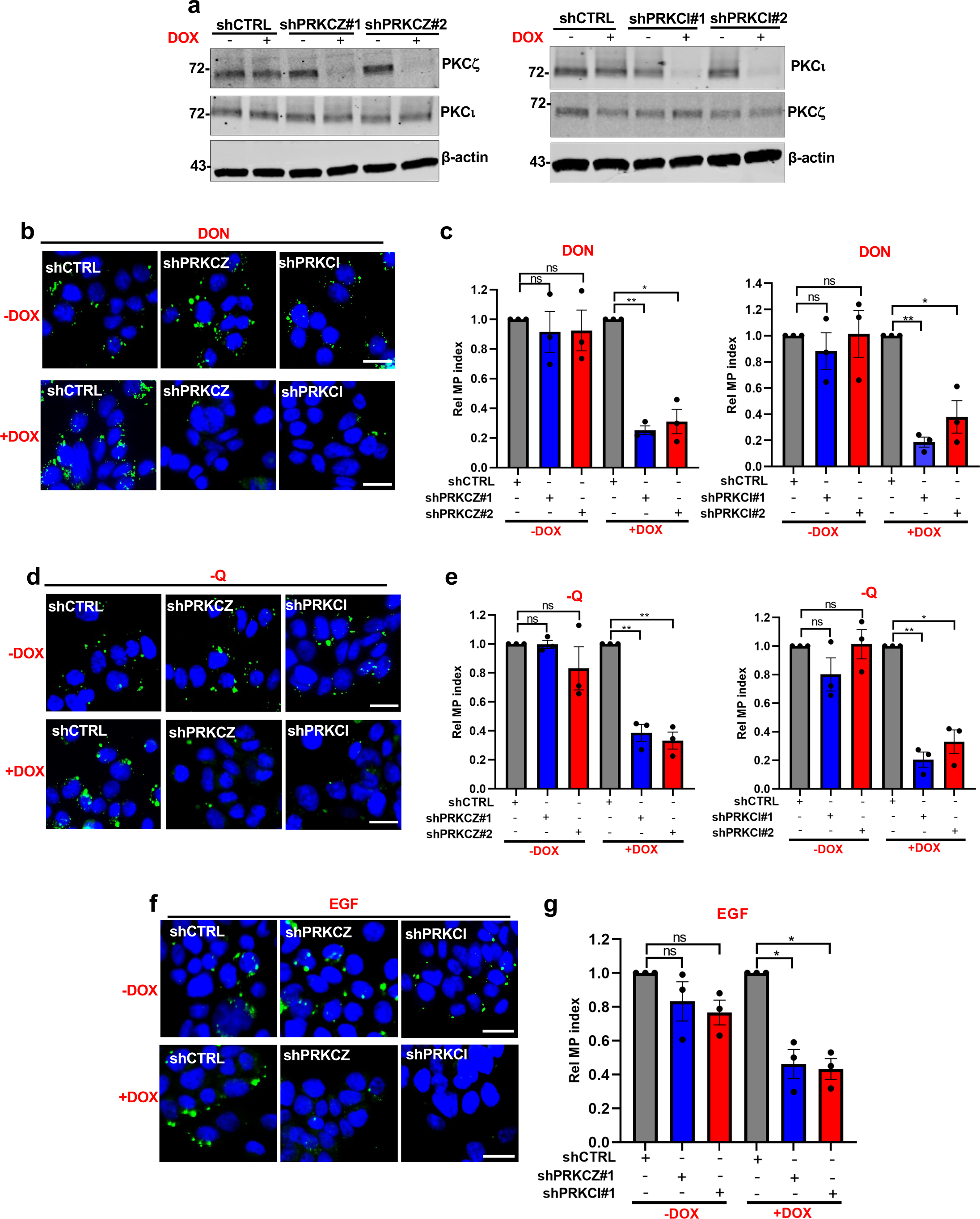
Assessment of macropinocytosis in AsPC-1 cells transduced with doxycycline-inducible shRNA hairpins targeting *PRKCZ* or *PRKCI*. **a)** Immunoblots assessing total PKCζ and PKCι protein levels in AsPC-1 cells transduced with either a non-targeting shCTRL, or one of two shRNAs targeting *PRKCZ* or *PRKCI*. Cells were cultured for 72 hrs with or without doxycycline (2 μg/ml). β-actin was used as a loading control. **b)** Representative images of macropinocytosis in DON-treated AsPC-1 cells cultured with or without doxycycline and transduced with shCTRL, shPRKCZ or shPRKCI. Scale bar, 20 μm. **c)** Quantification of macropinocytosis in DON-treated AsPC-1 cells cultured with or without doxycycline and transduced with the indicated shRNAs. Cells were cultured with or without doxycycline. Data are shown relative to the DON-treated shCTRL cells and are presented as the mean ± SEM of three independent experiments. **d)** Representative images of macropinocytosis in glutamine-deprived AsPC-1 cells cultured with or without doxycycline and transduced with shCTRL, shPRKCZ or shPRKCI. Scale bar, 20 μm. **e)** Quantification of macropinocytosis in glutamine-deprived AsPC-1 cells cultured with or without doxycycline and transduced with the indicated shRNAs. Cells were cultured with or without doxycycline. Data are shown relative to the glutamine-starved shCTRL cells and are presented as the mean ± SEM of three independent experiments. **f)** Representative images of macropinocytosis in EGF-treated (100 nM) AsPC-1 cultured with or without doxycycline and transduced with shCTRL, shPRKCZ or shPRKCI. EGF was added for 10 min prior to macropinocytosis analysis. Scale bar, 20 μm. **g)** Quantification of macropinocytosis for the conditions described in f. Data are shown relative to shCTRL cells and are presented as the mean ± SEM of three independent experiments. Statistical significance was calculated using unpaired two-tailed Student’s t test with Welch’s correction. ns, non-significant, \**P*<0.05, \*\**P*<0.01.

**Extended Data Fig. 8.**
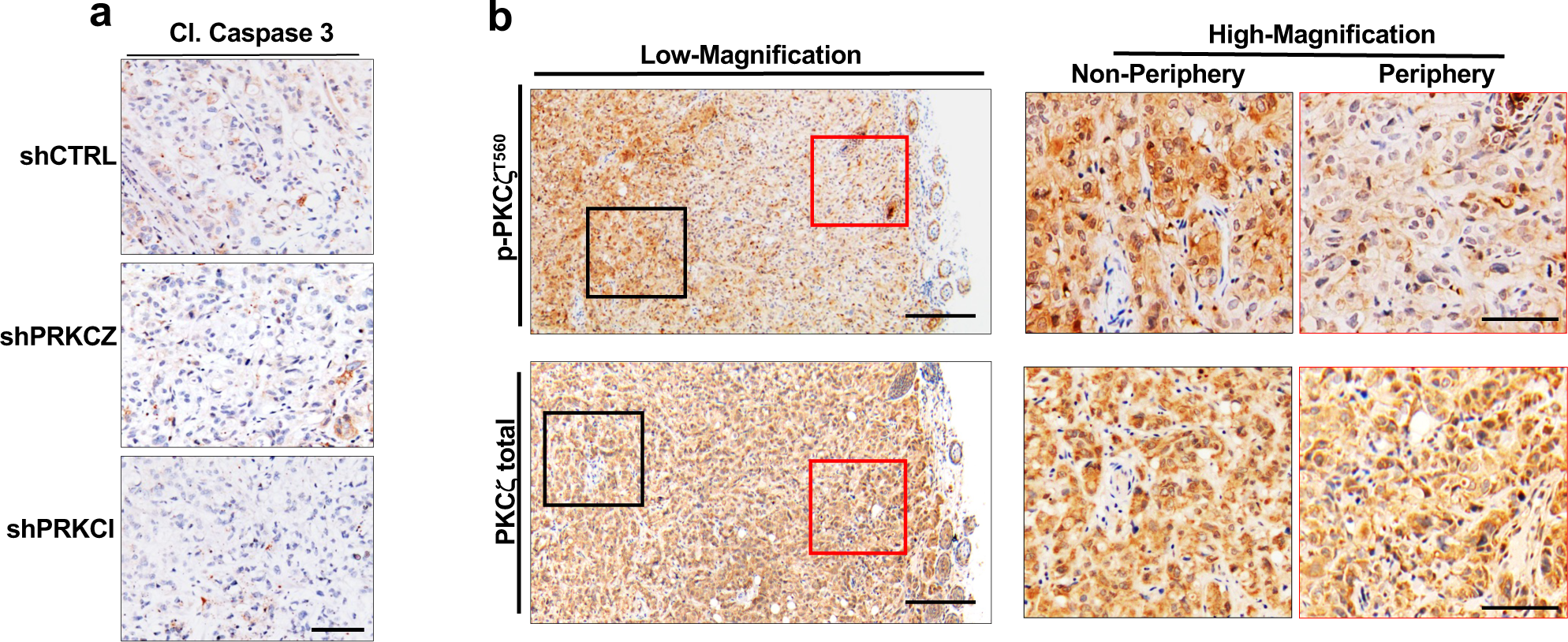
aPKC depletion does not affect apoptosis *in vivo* and PKCζ phosphorylation is enhanced in tumor cores. **a)** Immunohistochemical analysis of cleaved-caspase 3 (Cl. Caspase 3) in AsPC-1-derived xenograft tumors. Data are representative of n=4 different tumors per condition. Scale bar, 100 μm. **b)** Immunohistochemical analysis of p-PKCζ^T560^ and total PKCζ in AsPC-1 control xenografts. Data are representative of n=4 different tumors per condition. Scale bar, 100 μm. Images on the right represent a higher magnification of the boxed areas within the images on the left. Scale bar, 50 μm.

**Extended Data Fig. 9.**
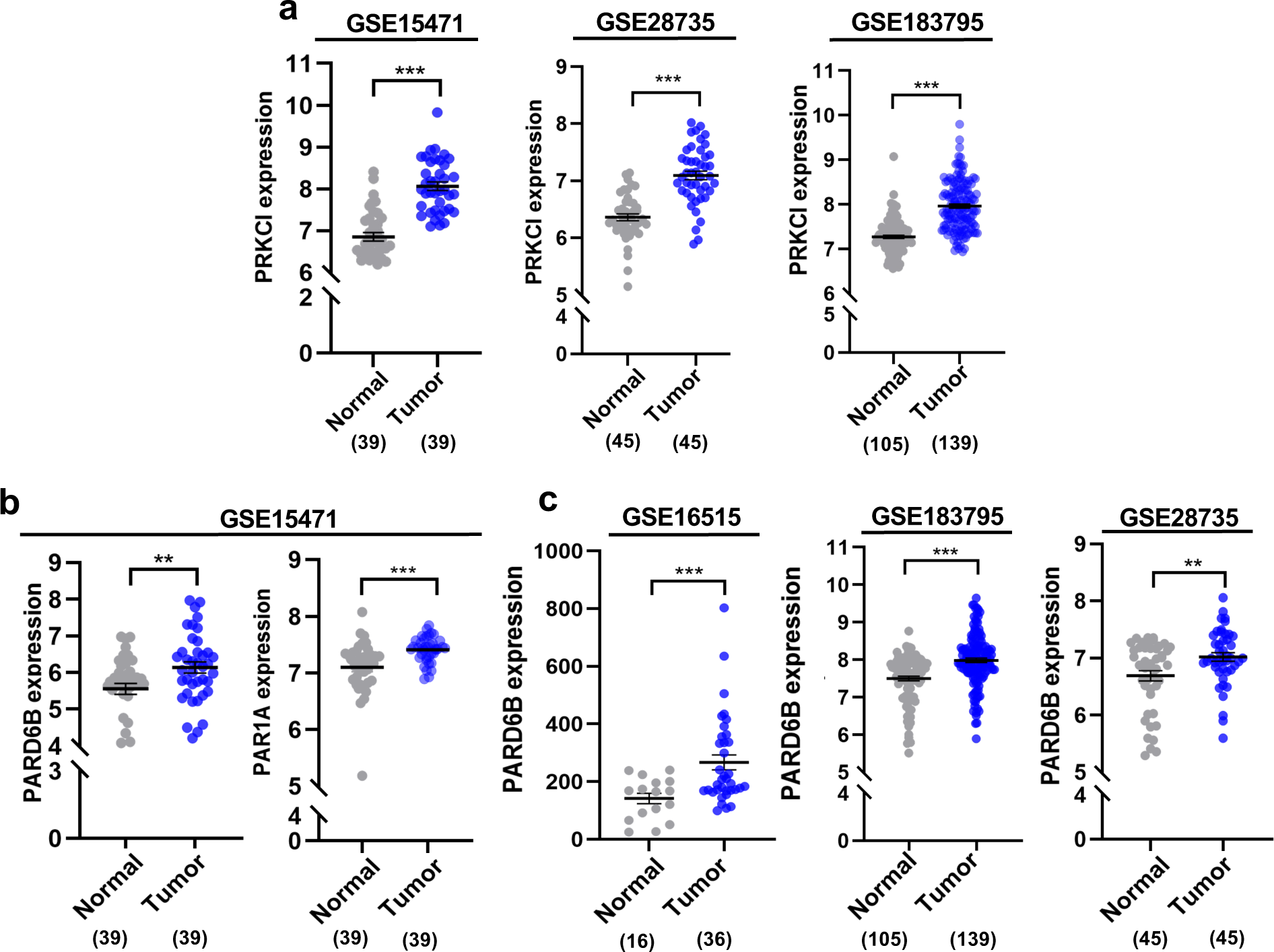
*PRKCI*, *PARD6B* and *PAR1A* expression in PDAC patient datasets. **a)** *PRKCI* transcript levels in normal tissue versus PDAC from the indicated GSE dataset. Sample numbers per condition are indicated in parentheses. **b)** *PARD6B* and *PAR1A* transcript levels in normal tissue versus PDAC from GSE15471. Sample numbers per condition are indicated in parentheses. **c)** *PARD6B* transcript levels in normal tissue versus PDAC from the indicated GSE dataset. Sample numbers per condition are indicated in parentheses. Statistical significance was calculated using unpaired two-tailed Student’s t test. \*\**P*<0.01, \*\*\**P*<0.001.

